# Large-scale phylogenomics uncovers a complex evolutionary history and extensive ancestral gene flow in an African primate radiation

**DOI:** 10.1101/2023.06.21.545890

**Authors:** Axel Jensen, Frances Swift, Dorien de Vries, Robin Beck, Lukas F.K. Kuderna, Sascha Knauf, Idrissa S. Chuma, Julius D. Keyyu, Andrew C. Kitchener, Kyle Farh, Jeffrey Rogers, Tomas Marques-Bonet, Kate M. Detwiler, Christian Roos, Katerina Guschanski

## Abstract

Understanding the drivers of speciation is fundamental in evolutionary biology, and recent studies highlight hybridization as a potential facilitator of adaptive radiations. Using whole-genome sequencing data from 22 species of guenons (tribe Cercopithecini), one of the world’s largest primate radiations, we show that rampant gene flow characterizes their evolutionary history, and identify ancient hybridization across deeply divergent lineages differing in ecology, morphology and karyotypes. Lineages experiencing gene flow tend to be more species-rich than non-admixed lineages. Mitochondrial transfers between distant lineages were likely facilitated by co-introgression of co-adapted nuclear variants. Although the genomic landscapes of introgression were largely lineage specific, we found that genes with immune functions were overrepresented in introgressing regions, in line with adaptive introgression, whereas genes involved in pigmentation and morphology may contribute to reproductive isolation. This study provides important insights into the prevalence, role and outcomes of ancestral hybridization in a large mammalian radiation.

## Introduction

Ancient hybridization has been reported in many organisms, including mammals (Taylor and Larson 2019). However, owing to their large genomes, studies of entire mammalian radiations, particularly among species-rich groups, are underrepresented (but see (Gopalakrishnan et al. 2018; Chavez et al. 2022), and most cases focus on pairs of species (Taylor and Larson 2019). Yet, large radiations with lineages of different ages, offer a unique opportunity to understand how increasing lineage divergence along a speciation continuum may impact the ability to hybridize and the outcome of hybridization events. As reproductive isolation accumulates with genetic distance (Coyne and Orr 2004), the impact and consequences of introgression may vary, as well as the mechanisms that allow for genomic incompatibilities to be overcome.

Several studies have reported that introgression between divergent lineages may spark rapid speciation and adaptive radiations by introducing novel genetic variation (Marques et al. 2019; Meier et al. 2017; Svardal et al. 2020). If so, we should expect a general pattern of elevated species richness in clades that have experienced ancient hybridization with evolutionary distinct lineages, whereas unadmixed lineages should contain fewer species. This prediction can only be studied with comprehensive genomic datasets from species-rich radiations.

Here we focus on guenons (tribe Cercopithecini), a species-rich group of African primates that radiated over the last ca. 10-15 million years (Guschanski et al. 2013; Kuderna et al. 2023). With 89 taxa, of which more than 30 are currently recognized as distinct species (IUCN 2022), guenons represent one of the world’s largest primate radiations. They provide a good case study of the speciation continuum, as multiple clades radiated independently during the last ca. 12 million years (Kuderna et al. 2023) and thus span a broad range of divergence times. Guenons are renowned for their ecological, morphological and karyotypic diversity and have attracted the attention of evolutionary biologists and ecologists for decades (Grubb et al. 2003; Campbell et al. 2007; Moulin et al. 2008; Glenn and Cords 2002; Gautier-Hion et al. 1988). Yet, despite possessing multiple characteristics that act as reproductive barriers in other study systems, guenons readily hybridize, producing viable and at least partially fertile offspring (Detwiler et al. 2005; de Jong and Butynski 2010; Detwiler 2019). Genomic studies have also identified ancient gene flow among a few guenon lineages (van der Valk et al. 2019; Ayoola et al. 2020; Svardal et al. 2017), but the extent and role of ancestral hybridization throughout the clade have not been investigated.

As such, guenons provide a highly informative study system to i) explore whether ancient hybridization between divergent lineages leads to greater species diversity, ii) study genomic patterns of introgression between lineages that differ in evolutionary divergence and uncover mechanisms that allow barriers to gene flow to be overcome at different evolutionary distances, iii) investigate the repeatability of introgression landscapes along the speciation continuum, and infer the predictability of genomic loci putatively involved in reproductive isolation, iv) explore the functional role of introgressed regions and their potential to contribute to lineage-specific adaptations, and v) test for the role of karyotypic differences in speciation.

## Results

### Dataset, Sequencing and Genotyping

We compiled a dataset of high coverage, whole genome sequences from 37 samples belonging to 24 primate species ((Kuderna et al. 2023; van der Valk et al. 2019; Ayoola et al. 2020), Table S1). These included 22 guenon species from all six genera, *Allenopithecus, Allochrocebus, Chlorocebus, Erythrocebus, Miopithecus* and *Cercopithecus* (Figure 1A), which collectively show a high degree of sympatry throughout sub-Saharan Africa (Figure 1B). The most species-rich genus, *Cercopithecus*, is commonly divided into six evolutionary distinct species groups: *cephus*, *mitis*, *mona*, *neglectus*, *diana* and *hamlyni* (Grubb et al. 2003; Lo Bianco et al. 2017), all of which are represented in our dataset. We also included two outgroup species from the sister tribe Papionini: *Macaca mulatta* and *Cercocebus torquatus*. Average read mapping depth varied between 15.7 and 57.7, with a median of 29.8. After calling genotypes against the rhesus macaque reference genome (Mmul_10, GenBank: GCA_014858485.1) and applying a stringent set of filters (Methods), we obtained a total of 1.18 billion genotyped sites across the autosomes, 52 million sites on the X-chromosome and 63,565 sites on the Y-chromosome. Of these, 140 and 4.2 million sites were bi-allelic SNPs on the autosomes and X-chromosome, respectively. Additionally, we assembled and annotated the mitochondrial (mtDNA) genomes of all samples.

**Figure 1.**
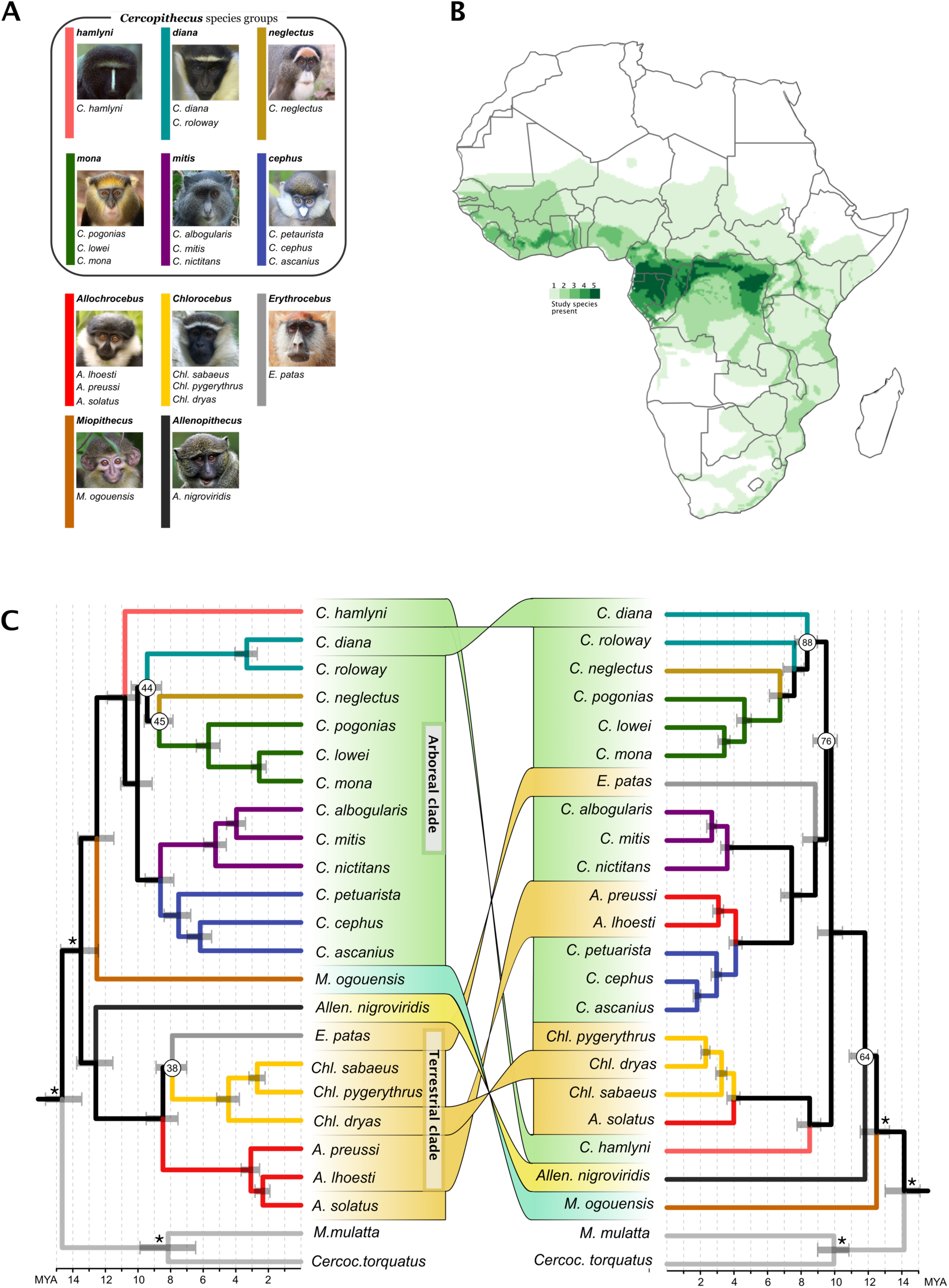
Taxonomy, species richness and mito-nuclear discordances among guenons. A) Taxonomic overview of the 22 species included in this study, shown according to genus and species groups. The color of the vertical bars corresponds to branch colors in (C). B) Species richness/degree of sympatry for species included in this study, based on species distributions from IUCN (IUCN 2022) C) Astral species tree obtained from 3,346 autosomal gene trees (left) and maximum likelihood tree constructed with RaxML from complete mitochondrial genomes (right), with connectors highlighting phylogenetic discordances. Node annotations show branch quartet support (left) or bootstrap support (right), where this was less than 95 %. Branches are colored based on genus/species groups as in (A). Tree topologies were estimated from all available samples (Figure S1, S7) and were subsequently pruned to a single sample per species prior to divergence date estimates with MCMCTree, applying fossil calibrations to the nodes annotated with asterisks. Nodes were rotated to aid visualization of mito-nuclear discordances. Photo credit for A): *hamlyni*: NRowe/alltheworldsprimates.org; *diana*, *Chlorocebus*: K. Guschanski; *neglectus*: M. D’haen; *mona*: S. Knauf; *mitis*: M. Mpongi & K. Detwiler; *cephus*: S. Crawford & K. Detwiler; *Allochrocebus*: T. <u>Ukizintambara;</u> *Erythrocebus*: T. Valkenburg; *Miopithecus*: P. Paixão; *Allenopithecus*: D. Sutherland.

### Species tree inferences

We constructed a multi-species coalescence tree in ASTRAL (Zhang et al. 2018), using 3,346 independent autosomal gene trees, each representing a 25 kb genomic region sampled every 500 kb along the genome (Figure 1C, S1, Methods). All genera, species groups and species were monophyletic with maximal local posterior probability (lpp) support (lpp=1). The tree topology was largely consistent with that recovered in a large primate phylogenetic analysis based on ca. 3,500 ultra-conserved element (UCE) loci (Kuderna et al. 2023) except for the placement of the genus *Erythrocebus* and the *diana*-species group. Both these discrepancies involve rapid diversifications with short internal branches and strong incomplete lineage sorting (ILS), as illustrated by high degree of gene tree discordance around these nodes (normalized quartet score = 38 % and 44 %, respectively). The slow evolution of UCE loci (Bejerano et al. 2004) may reduce their power to resolve rapid radiations, potentially explaining these differences.

We estimated divergence times on the ASTRAL topology using four independent MCMCTree runs (Yang 2007). After confirming that the runs converged at highly similar node age estimates, they were merged and analyzed jointly, resulting in an effective sample size (ESS) ≥ 285 for all nodes (Figure S2). These analyses suggest that the guenons split from Papionini ca. 14.6 million years ago (MYA, Figure 1C, Table S2), followed by the first split within the guenons, dividing them into two major clades ca. 13.4 MYA. One of these clades contains the genera *Miopithecus* (*M*) and *Cercopithecus* (*C*), which diverged from each other ca. 12.4 MYA. The mainly arboreal species groups that make up the genus *Cercopithecus* radiated ca. 8-11 MYA and will hereafter be referred to as the ’Arboreal clade’. The other major clade contains the genus *Allenopithecus* (*Allen*), as sister to the terrestrial genera *Chlorocebus* (*Chl*), *Erythrocebus* (*E*) and *Allochrocebus* (*A*), hereafter termed the ’Terrestrial clade’. The Terrestrial clade split from *Allenopithecus* ca. 12.5 MYA and underwent rapid radiation ca. 8 MYA, forming the three terrestrial genera.

We repeated the same process with 118 X-chromosomal loci, resulting in an identical topology with highly similar divergence date estimates (Figure S3-S5). The placement of *Erythrocebus* as a sister to *Chlorocebus* received low support (lpp = 0.53), likely due to rapid radiations and a high degree of incomplete lineage sorting (ILS).

Samples from male individuals were available for 17 guenon species and were used to call Y-chromosomal genotypes. These sites were concatenated into a multi-sequence alignment, and evolutionary relationships were inferred with RaxML (Stamatakis 2014). Compared to the autosomal tree, the Y-chromosomal tree disagreed only in the placement of *Erythrocebus*, in line with the incomplete support obtained for this node in other datasets (Figure S6). Considering the high topological concordance among nuclear markers of different inheritance modes, we hereafter refer to the autosomal phylogeny as the ’species tree’ (Figure 1C), although some uncertainty remains regarding the exact placement of *Erythrocebus*.

### Mito-nuclear discordances suggest ancient hybridization

We generated assemblies for the mitochondrial genomes of all study species and used these to construct a maximum likelihood tree in RaxML, and estimated divergence dates with MCMCTree (Figure 1C, S7, S8, Methods). The mitochondrial phylogeny was consistent with that previously recovered from a large dataset of museum specimens (Guschanski et al. 2013) and showed several hard incongruences with the species tree. In the mitochondrial tree *Miopithecus* is sister to all guenons, followed by *Allenopithecus* as sister to all Arboreal and Terrestrial clades members, from which they diverged 11-12 MYA. We also found several mito-nuclear incongruences among the Arboreal and Terrestrial clades: The terrestrial genus *Erythrocebus* is nested within the Arboreal clade, sister to the *cephus* and *mitis* species groups, and two terrestrial *Allochrocebus* species – *A*. *lhoesti* and *A*. *preussi* – cluster together with the arboreal *cephus* species group, whereas their congener *A. solatus* is sister to the terrestrial genus *Chlorocebus*. Additionally, we found several discordances within genera, confirming previous reports. In contrast to the species tree, *Chl. dryas* is sister to *Chl. pygerythrus* ((van der Valk et al. 2020), Figure 1C), and *C. mona* is paraphyletic, as the sample from the eastern population clusters with *C. pogonias* ((Ayoola et al. 2020) Figure S7). The *diana* species group is also paraphyletic, in agreement with previous mtDNA genome studies (Guschanski et al. 2013).

Topological differences between nuclear and mitochondrial phylogenies are not uncommon and can be caused by ILS or hybridization. However, for ILS to generate conflicting topologies in which a lineage changes position from being nested in one clade to being nested within another, the mitochondrial genome must remain polymorphic in several lineages throughout multiple speciation events, which is unlikely. Thus, ILS remains a plausible explanation for the conflicting positions of, e.g., *Miopithecus* and *C. diana*, for which the retention of mitochondrial polymorphism is required only across a single split. In contrast, most mito-nuclear discordances detected here are more likely explained by mitochondrial introgression, suggesting several cases of ancient hybridization event.

### Ancient gene flow is prevalent among guenons

To further explore the possibility of ancient hybridization events, we used Dsuite (Malinsky et al. 2021) to calculate the D-statistics and f4-ratios on autosomal data (Durand et al. 2011) for all possible trios of guenon species, using the papionin *Macaca mulatta* (Rhesus macaque) as outgroup. Out of 1,540 tested trios, 865 produced significantly positive D-statistics, indicative of gene flow. However, these D-values may be interdependent, resulting either from ancestral gene flow events, creating similar D-values in all descendants, or from the introduction of shared ancestral alleles, causing significant D-statistics in sister species that did not exchange genes. To disentangle and polarize this complex pattern, we used different combinations of D-statistics similar to the dfoil-method (Pease and Hahn 2015) together with an approach adapted from the partitioned D-statistic (Eaton and Ree 2013), in which we excluded shared variants to detect ’carryover’-effects (see Methods). Such carryover effects may produce a false positive signal of gene flow in lineages that are close relatives to the actual source as a consequence of their shared ancestry. Identifying these effects allowed us to disentangle the primary source and recipient lineages (i.e., infer directionality) for most gene flow events.

We detected the strongest signal of excess allele sharing, indicative of ancestral hybridization, between the *cephus* and *mona* species groups (D = 0.15-0.29, Figure 2, events A1 and A2 in Figure 3). The f4-ratios, which provide an estimate of the proportion of introgression across the genome, suggested that at least 10-20 % of the genome was introgressed (Svardal et al. (2020) found through simulations that f4-ratios may substantially underestimate the actual proportion of gene flow). This gene flow event also produced significant values of D between the *cephus* group and *C. neglectus*, the latter being sister to the *mona* group (Figure 2). However, this was likely driven by shared ancestral variation between *C. neglectus* and the *mona* group, as these D-values approached zero when only private alleles were considered (Figure 2). This suggests directional gene flow from the *mona* group, with carryover effects from *C. neglectus*, into the *cephus* group.

**Figure 2.**
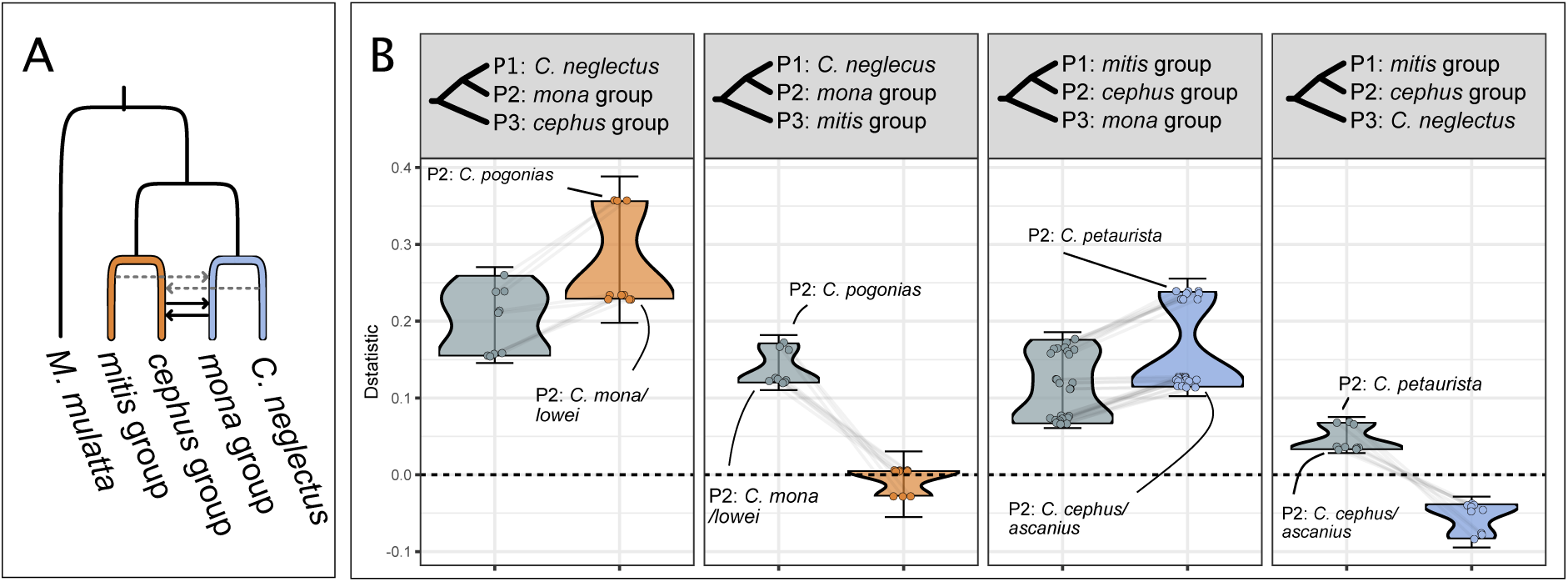
D-statistics and partitioned D-statistics using gene flow event A (Figure 3) as an example. A) Schematic overview of gene flow event A (Figure 3), with branch colors corresponding to the partitions used to estimate private allele sharing. Solid black arrows show the inferred gene flow, and dashed grey arrows show allele sharing inferred as ‘carry-over’ effects. B) Four tests for excess allele sharing between the *mona* group, *cephus* group, *mitis* group and *C. neglectus*. Grey points and distributions show the D-statistics for all combinations of taxa from the respective groups as depicted in the panel header, with *M. mulatta* as outgroup, using all SNPs. Colored points and violin distributions show the D-statistics for the same trios (connected by grey lines) after removing sites with shared alleles between the *cephus* and *mitis* groups (orange) or *C. neglecus* and the *mona group* (blue). Error bars show the lowest and highest D-statistic +/-3 standard errors. When D-statistics approach zero after removing shared alleles, we interpret this as carryover effects, indicating directionality. Note that some D values turn negative after removing shared ancestral variation, which is an expected consequence of this method and should not be interpreted as excess allele sharing between P1 and P3 (Pease and Hahn 2015).

**Figure 3.**
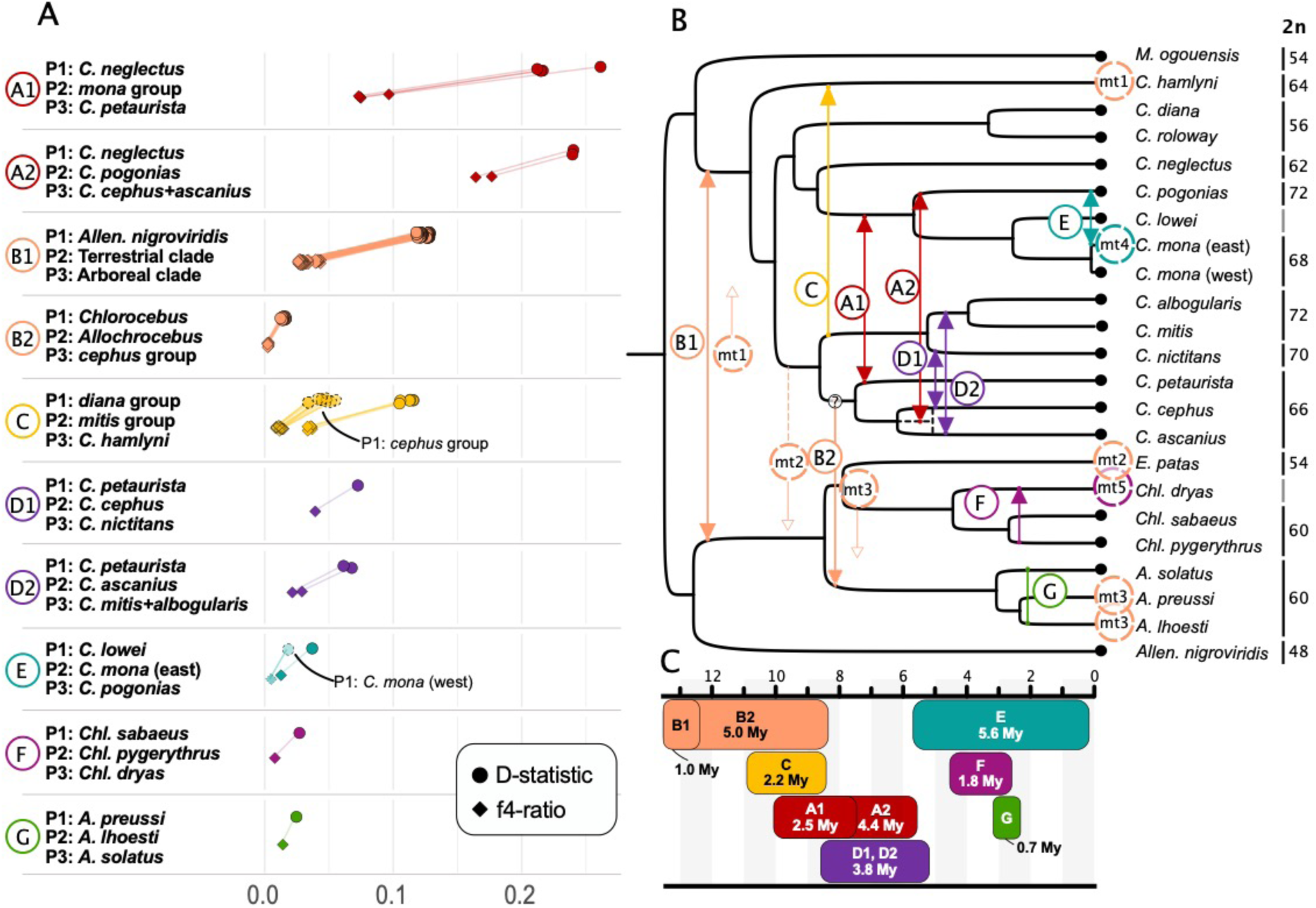
Excess allele sharing and gene flow events among guenons. A) D-statistic (circles) and f4-ratios (diamonds) as estimated by Dsuite illustrating excess of allele sharing caused by the seven identified gene flow events. The tested taxa combinations are shown on the left (with *M. mulatta* as outgroup). Positive D-statistic values indicate gene flow between P3 and P2 taxa and are connected by lines to the f4-ratios that reflect the proportion of gene flow. The values were obtained for all possible members of the respective species groups, for which more than a single representative was available in our dataset. In events C and E, we observed decreasing D-statistics with more closely related P1 taxa (annotated as dashed shapes), indicating directionality from P2 into P3. B) Schematic overview of the identified gene flow events. Arrows are shown where directionality could be inferred and mitochondrial introgressions, accompanying nuclear gene flow between internal branches, are shown as dashed circles with an arrow. Dashed circles on tips highlight species that carry an introgressed mitochondrion, with circle color corresponding to the most likely introgression event. The donor in B2 was not fully resolved, as illustrated by the question mark, and the dashed line of the *C. cephus*/*ascanius* divergence illustrates the likely overestimate of this node’s age (see main text). Where available, annotations to the right of species labels show the chromosome number of extant species. C) Schematic overview of the minimum time since divergence at the time of introgression, based on the divergence time estimates. The divergence time estimate between the eastern and western population of *C. mona* was retrieved from Ayoola et al. (2021).

Similarly, we found excess allele sharing between the *mona* and *mitis* groups, the latter being sister to the *cephus* group, also primarily driven by carryover effects from shared ancestry between the *cephus* and *mitis* groups (Figure 2). Hence, we conclude that there has been bidirectional gene flow between the ancestral lineages of the *cephus* and *mona* groups. Within the *cephus* and *mona* groups, the D-values varied substantially across species (Figure 2), suggesting more than a single ancestral pulse of gene flow. The simplest scenario that could explain the observed D-values involves two independent bidirectional gene flow events: one between *C. petaurista* and some members of the *mona* group, most likely their common ancestor, and a second between *C. pogonias* and the ancestor of *C. cephus*/*ascanius* (A1 and A2 in Figure 3). Notably, the split between *C. cephus* and *C. ascanius* pre-dates that between *C. pogonias* and *C. mona/lowei* in our divergence time estimates, which conflicts the suggested second pulse into their common ancestor. However, divergence time estimates from genomic regions with low levels of introgression between the *cephus* and *mona* groups produced a substantially younger split time between *C. cephus* and *C. ascanius*, post-dating the divergence between *C. pogonias* and *C.* mona/*lowei* by ca. 1 million years (MY) (Figure S9). These estimates are broadly in line with the divergence times proposed by (Kuderna et al. 2023) using UCE loci and suggest that our estimates for the *cephus* group are likely inflated due to different levels of introgression from the *mona* group.

The second strongest signal of excess allele sharing was detected between the Arboreal and Terrestrial clades (D ∼0.12, f4-ratio ∼ 0.03-0.04, event B1 and B2 in Figure 3). Using the same polarization approach, we found that most of the observed D-values can be attributed to a bidirectional gene flow event between the ancestors of the two clades, ca. 11-12 MYA (event B1, Figure S10). However, we also found that the terrestrial genus *Allochrocebus* shares significantly more alleles with the Arboreal clade than its sister genus *Chlorocebus* (Figure S11A), suggesting that gene flow between the Arboreal and the Terrestrial clades continued or reoccurred, along the ancestral *Allochrocebus* branch (event B2, Figure 3). This scenario is also supported by the presence of a *cephus* group-like mtDNA in *A. lhoesti*/*preussi* (Figure 1C, discussed below) and a trend towards excess allele sharing between specifically the *cephus* group and *Allochrocebus* (Figure S11B). The reason that this trend is largely not significant could be due to the extensive gene flow within the Arboreal clade (Figure 3).

Furthermore, we find evidence for gene flow between the ancestor of the *mitis* group and *C. hamlyni* (Figure 3, event C, D ∼ 0.11, f4-ratio = 0.04), the *cephus* and *mitis* groups (event D, two pulses of gene flow, D ∼ 0.07, f4-ratio = 0.02-0.04, Figure 3, S12), *C. pogonias* and the eastern population of *C. mona* (event E, D = 0.04, f4-ratio = 0.01; also reported by (Ayoola et al. 2020), *Chl. dryas* and *Chl. pygerythrus* (event F, D = 0.03, f4-ratio = 0.01; as reported by (van der Valk et al. 2020) and *A. solatus* and *A. lhoesti* (event G, D = 0.02, f4-ratio = 0.01). Using the polarization approach, we could infer that gene flow was bidirectional in event D. Based on increasing D-statistics with more distant P1 lineages, it is most parsimonious to infer directional gene flow from the *mitis* group into *C. hamlyni* (event C) and from the eastern *C. mona* populations into *C. pogonias* (event E). However, we cannot test for the opposite direction of gene flow in these two events since our dataset lacks sister lineages to *C. hamlyni* and *C. pogonias*.

In total we pinpoint at least seven major ancestral gene flow events (A-G, Figure 3), some of which occurred in several pulses (A1 and A2, B1 and B2, D1 and D2, treated as separate pulses of the same event as they are strongly interdependent). However, we fully acknowledge that the presented scenarios are likely an oversimplification, and additional gene flow events may have occurred. By calculating the difference in age between the nodes separating P2 from its closest sister and that separating P3 from P2, we retrieved a minimum time of divergence at the time of gene flow for each event. These times ranged from 1 to over 5 million years (Figure 3C), suggesting that reproductive isolation may remain incomplete for long periods. To test the support for the inferred hybridization events with an independent, model-based approach, we also constructed phylogenetic networks in PhyloNetworks (Solís-Lemus et al. 2017; Solís-Lemus and Ané 2016). PhyloNetworks does not allow multiple reticulations along the same branch and is thus restricted to less detailed inferences. However, when analyzing subsets of the tree to test each independent event, we found strong support for all of them, with at least one pulse per event being identified (Figure S13-S20).

Among the identified gene flow events, events D and E most likely involved lineages with different chromosome numbers based on the known karyotypes of the extant species (Sineo et al. 1986; Lo Bianco et al. 2017)(Figure 3). To investigate if this was the case in older events, we reconstructed the ancestral karyotypes using ChromEvol (Höhna et al. 2016; Freyman and Höhna 2018), using different probabilities of chromosomal fission events. Each reconstruction returned unique ancestral states and converged poorly, resulting in low support values (lpp ≤ 25 for internal nodes with differences in chromosome numbers among descendants, Figure S21-22). Guenon karyotypes are highly homoplastic, possibly affected by hybridization (Moulin et al. 2008), which likely contributed to the difficulties in reconstructing the ancestral states. However, all reconstructions suggest that the more ancient events A (primarily A2), B, and C occurred across different karyotypes. Thus, five out of seven major gene flow events are most likely examples of cross-karyotypic gene flow. The remaining two events occurred within the karyotypically stable *Allochrocebus* and *Chlorocebus* genera (F and G), although the karyotype of *Chl. dryas* is not known.

### A combination of introgression and incomplete lineage sorting best explains mito-nuclear discordances

The detected gene flow events can directly explain three of the six mito-nuclear discordances (Figure 1C, Figure 3, events B, F, and E), whereas the remaining three cases are more complex. Our results suggest that these discordances are best explained by a combination of introgression and ILS, with mitochondria being introgressed ancestrally in a polymorphic state and subsequently retained and fixed only in a subset of the descending lineages. For example, in the mitochondrial phylogeny the Arboreal and Terrestrial clades are sisters, compatible with mitochondrial introgression in event B. However, the difference in the position of the *hamlyni* group, which is the sister of the rest of the Arboreal clade in the nuclear phylogeny, but placed as a sister to the Terrestrial clade in the mitochondrial phylogeny, is not supported by additional gene flow between *hamlyni* and the Terrestrial clade (Figure S23). A scenario in which a Terrestrial-like mtDNA was introgressed into the Arboreal clade ancestor during event B1 and subsequently retained only in the *hamlyni* group is thus most compatible with our results (Figure 3B). Similarly, despite indistinguishable levels of nuclear gene flow between all members of the genus *Allochrocebus* and the Arboreal clade (Figure S24), only the *A. lhoesti/*preussi lineage shows the introgressed *cephus* group-like mtDNA. This haplotype was most likely transferred into the common *Allochrocebus* ancestor in event B2 but differentially retained during subsequent divergence. The Arboreal-like mtDNA genome of the genus *Erythrocebus* is also likely the result of ancestral introgression and subsequent differential sorting, as this lineage shows lower levels of nuclear gene flow with the Arboreal clade than other members of the Terrestrial clade (Figure S10). A complicating factor is that this scenario requires the retention of mitochondrial polymorphism over two speciation events according to our species tree. However, as mentioned above, uncertainty remains regarding as to the precise phylogenetic position of *Erythrocebus*, as alternative topologies place it as a sister to a clade containing the *Chlorocebus* and *Allochrocebus* genera (Kuderna et al. 2023). Given this latter placement, the mtDNA polymorphism only needs to be retained over a single speciation event.

### Mitochondrial introgression across deeply divergent lineages was facilitated by the co-sorting of alleles in nuclear genes with mito-nuclear interacting functions

We identified several mitochondrial introgressions across deeply divergent lineages, combining nuclear and mitochondrial genomes that evolved independently for extended time periods. Nuclear and mitochondrial genes interact with each other during the essential production of ATP via oxidative phosphorylation (OXPHOS), and several reports attributed hybrid inviability to malfunctioning mito-nuclear complexes (reviewed by (Burton 2022)). We therefore investigated whether mitochondrial introgressions were facilitated by co-introgression of mito-nuclear interacting genes. We tested this in one of the most extreme cases, the introgression of a *cephus* group-like mitochondrion into the ancestral *Allochrocebus* lineage, involving lineages separated by at least five million years of independent evolution (Figure 3C). While *A. lhoesti* and *A. preussi* are fixed for an introgressed mitochondrial genome, their sister, *A. solatus*, retains the ancestral mtDNA. To understand if this differential retention of mtDNA was facilitated by differential sorting of introgressed, co-adapted nuclear variation, we investigated if *A. lhoesti/preussi* were, on average, more *cephus*-like in nuclear genes known to interact with mitochondrial genes (N-mt genes, Table S3) compared to a control set of genes without such interactions. This would suggest broad-scale co-evolution between N-mt genes and the mtDNA as a mechanism facilitating mitochondrial transfers. However, N-mt and control genes showed similar levels of *cephus* group ancestry in *A. lhoesti/preussi*, and there was no difference in absolute divergence to *A. solatus*, suggesting a lack of broad-scale co-introgression of N-mt genes (Figure S26). Next, we assessed individual SNP patterns on a gene-by-gene basis to identify signatures of fine-scale co-evolution between the introgressed mtDNA and N-mt genes. Using the same sets of genes (N-mt and control), we quantified the amount of *cephus* group alleles retained only in either *A. lhoesti/preussi* or *A. solatus*. We used *Chlorocebus* to polarize the ancestral Terrestrial clade allele and counted two SNP patterns: category 1 – grouping *A. lhoesti*/*preussi* with the *cephus* group and *A. solatus* with *Chlorocebus* (in agreement with the mtDNA topology) and category 2 – grouping *A. solatus* with the *cephus* group and *A. lhoesti*/*preussi* with *Chlorocebus* (opposing mtDNA topology). If only ILS is involved, we expect both these SNP categories to be equally frequent. However, in the presence of a non-neutral process that favors the retention of co-adapted mito-nuclear variants, we expect an excess of category 1 SNPs. In line with co-adapted N-mt alleles accompanying the mitochondrial introgression in *A. lhoesti/preussi*, there was a clear excess of category 1 SNPs in N-mt compared to control genes (Figure 4, Table S4). We found 196 category 1 SNPs in N-mt genes and a maximum of 133 in the control gene sets, compared to 96 category 2 SNPs in N-mt and ≤ 131 in control genes. More than half of the category 1 N-mt SNPs were found in the genes *NDUFA10* and *LRPPRC* (70 and 31, respectively) and included several non-synonymous variants (three and one, respectively, Table S4). No non-synonymous *cephus*-like variants were found in *A. solatus* in these genes, and across all considered N-mt genes, we detected a greater number of *cephus*-like non-synonymous variants in *lhoesti/preussi* than in *solatus* (11 versus 1). Mutations in *NDUFA10* have been linked to mitochondrial complex 1 deficiency in humans (Hoefs et al. 2011), and mutations in *LRPPRC* have been associated with mitochondrial complex 4 deficiency (Oláhová et al. 2015), both of which are involved in oxidative phosphorylation. Our results thus suggest that retaining *cephus*-like variants in these genes may have been essential to overcome mito-nuclear incompatibilities and thus facilitated mitochondrial introgression.

**Figure 4.**
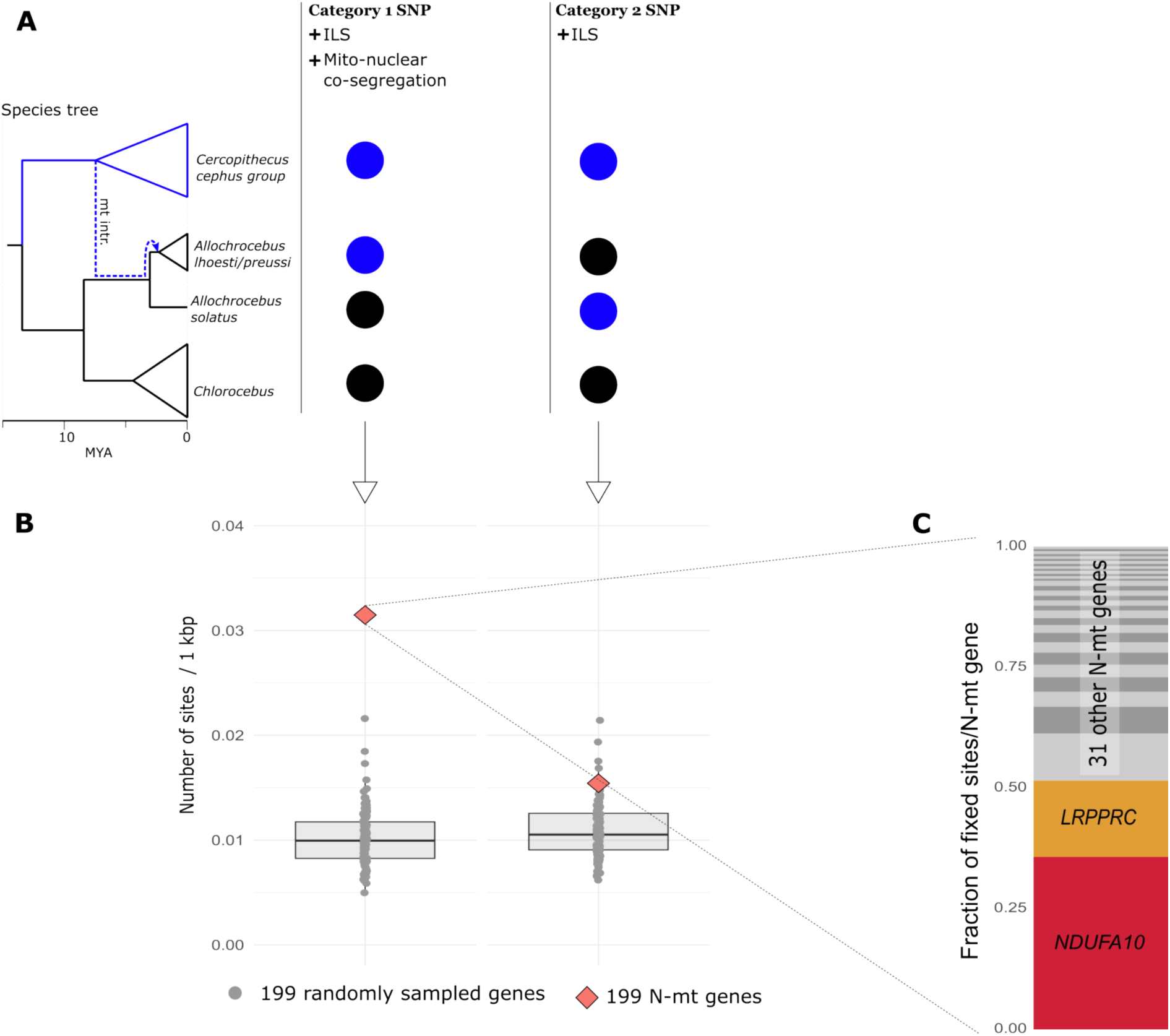
Prevalence of privately retained introgressed *cephus* group alleles in *A. lhoesti/preussi* and *A. solatus* in nuclear genes involved in mitochondrial functions (N-mt) compared to the genomic background. A) Schematic overview of the two considered SNP categories. The blue dashed line in the species tree illustrates the mitochondrial introgression from the *cephus* group to the ancestral *Allochrocebus* branch, subsequently retained only in the *A. lhoesti*/*preussi* lineage. The circles illustrate the two categories of SNPs differentially fixed between the *cephus* group (blue circles) and *Chlorocebus* (black circles): Category 1 groups *A. lhoesti/preussi* with *cephus* and can arise through ILS and mito-nuclear co-segregation, whereas category 2 groups *A. solatus* with *cephus* and is expected only from ILS. B) Number of SNPs per 1,000 base pairs of categories 1 and category 2 across 199 N-mt genes (red diamonds) and 100 samples of 199 other nuclear genes (grey dots). C) Proportion of category 1 SNP contributed by 33 N-mt genes with at least one such site, highlighting the identity of the two most important genes.

### Admixed genomes are a mosaic of different ancestries with elevated heterozygosity in introgressed regions

Next, we investigated the spatial patterns of gene flow to understand the genomic architecture of introgression. For simplicity, we only tested the trio setup ((P1,P2) P3) with the highest D-values for events occurring in several pulses (A1, B1, D1, Figure 3). We constructed neighbor joining trees in non-overlapping 25 kb windows along the genome and assigned them to three main topologies: Tree 1 corresponds to the species tree, Tree 2 shows the introgressed topology and Tree 3 is consistent with ILS. The distribution of the tree topologies across the genome revealed a highly heterogeneous landscape, with regions showing the species tree – the most frequent topology in all instances – being interspersed with short regions of introgressed and ILS-derived ancestry (Figure 5A, Figures S26-31). Consecutive windows (long segments) of introgressed ancestry were rare, in line with ancient gene flow where recombination acted for a long time to break up introgressed haplotype blocks (Figure S32). Notably, Tree 2 may also arise through ILS and the proportion of introgressed ancestry must be assessed relative to Tree 3, which is expected to result only from ILS. Tree 2 was always more prevalent than Tree 3 and, in line with the strength of gene flow inferred through D-statistics, we find the highest relative proportions of Tree 2 in events A-D.

**Figure 5.**
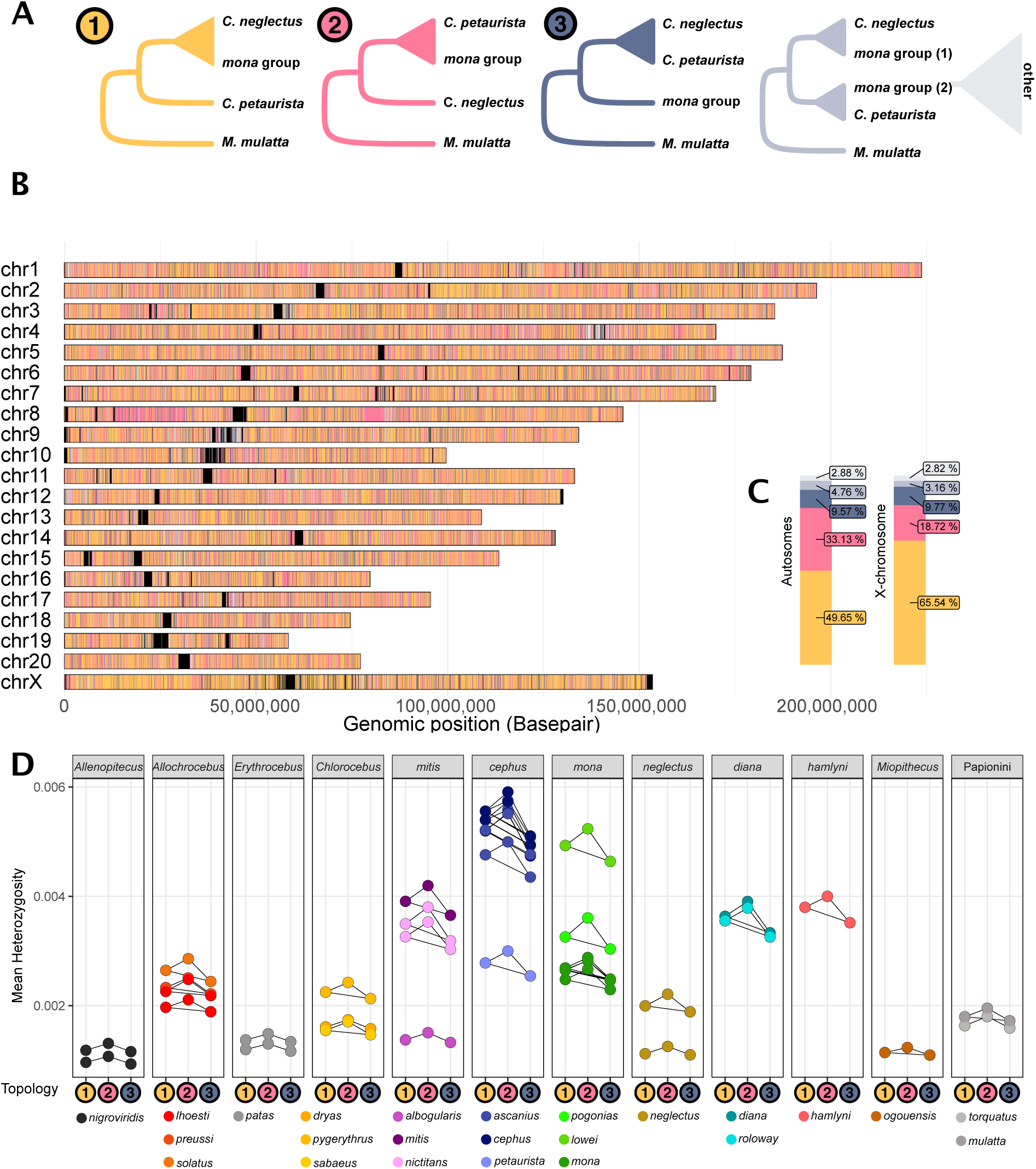
Distribution, prevalence and heterozygosity of introgressed genomic segments. A) Five possible tree topologies obtained from 25 kb genomic windows for the introgression event A1 (Figure 3), involving *C. neglectus*, *mona* and *cephus* groups (here represented by *C. petaurista*), rooted with *M. mulatta* as outgroup. Tree 1 groups the lineages according to the species tree. Tree 2 groups the *mona* and *C. petaurista* groups monophyletically to the exclusion of *C. neglectus* and can be caused by ancestral introgression or ILS. Tree 3 places the *cephus* group with *C. neglectus* and is expected to be caused only by ILS. The remaining two trees (light blue and gray) represent more complex topologies, which could be caused by, e.g. more recent introgression or ILS, respectively. B) The genomic location of the different tree topologies along the *M. mulatta* chromosomes, and C) their relative abundance on the autosomes and the X-chromosome. Black blocks in (B) correspond to regions of the genome with insufficient information for inferences, frequently located around centromeres and telomeres. D) Heterozygosity in windows corresponding to Tree 1, Tree 2, and Tree 3 topologies (shown as colored circles along the x-axis) calculated for all samples and species. Connectors show significant within-sample differences in heterozygosity between tree topologies, as assessed by ANOVAs followed by post hoc Tukey’s tests corrected for multiple testing.

To assess how genetic diversity relates to local ancestry, we calculated heterozygosity in the same 25 kb autosomal windows as above. Heterozygosity positively correlates with local recombination rate, mainly due to reduced linked selection (Nachman 2001; Cutter and Payseur 2013). The same mechanism is predicted to create a positive correlation between local introgression and recombination rate: introgressed haplotype blocks, which are predominantly deleterious, are efficiently purged in regions of low recombination, whereas high recombination rate may decouple neutral or beneficial alleles from their deleterious neighbors, leading to a higher local introgression rate (Veller et al. 2021; Figueiró et al. 2017). Without recombination maps for most of our study species, we used heterozygosity as a rough proxy for recombination rate, hypothesizing that it would be higher in regions with introgressed ancestry. Indeed, we observe this pattern in three of the four strongest gene flow events (A, B, and D), with Tree 2 windows showing higher heterozygosity than Tree 1 and Tree 3 windows (Figure 5D, S26D, S28D). Remarkably, this pattern holds for all taxa, not only those engaged in the particular gene flow event (*cephus* and *mona* in the event A, Figure 5D), and thus argues against higher heterozygosity being driven by segregating introgressed alleles, which would only affect the recipient lineages. Instead, it is consistent with a globally higher recombination rate in regions prone to retaining introgressed material. We do not find elevated heterozygosity in introgressed regions in some of the weaker gene flow events (C, E, F, G, Figure S27D, S28-S31D). It is possible that introgressed regions are too short and too few in these events or that the positive correlation between introgression and local recombination rate is not a universal pattern.

### Introgressed regions show low repeatability across independent gene flow events and are enriched for immune genes

Next, we quantified the introgression proportion in regions along the genome by calculating F_D_ in non-overlapping 25 kb windows (Martin et al. 2015). If genomic architecture and local recombination rate are the main determinants of the genomic location of introgressed regions and if synteny is conserved in different guenon species, we expect to repeatedly find the same genomic regions introgressing in separate gene flow events (Langdon et al. 2022). We found weak, albeit significant, correlations in the introgression landscape across all 21 comparisons (Pearson’s R^2^ range 0.008-0.100 p < 0.05, Figure S33), suggesting low levels of parallelism and repeatability.

We also investigated the functional relevance of introgression for each event, by identifying genes overlapping genomic regions with exceptionally high signals of introgression (F_D_ ≥ 99^th^percentile). To reduce the risk of false positives in this analysis, we called 25 kb window as introgressed only if they fulfilled a set of stringent criteria (Methods). We found 1,588 genes in highly introgressed regions (n = 188-345 per event, Table S5). Most of these genes were present in a single event (n = 1412), in line with the low repeatability of introgression. No significant enrichments of gene ontology (GO) categories were found when the gene lists from each event were tested separately (Fischer’s exact test corrected with false discovery rate [FDR] > 0.05). However, considering the entire set of introgressed genes, we detected an significantly enrichment of genes annotated with the GO-term ’regulation of lymphocyte mediated immunity (GO:0002706)’ (FDR = 0.0493, Table S6).

Albeit not present among functionally enriched categories, we note several introgressed genes involved in meiotic chromosome segregation (n = 11, Table S5) as they may play a role in cross-karyotypic gene flow events. Nine of these genes introgressed between lineages that most likely differed in karyotypes at the time of gene flow. Among them are SMC2, which is critical for bivalent chromosome formation during meiosis (Lee et al. 2011), and the well-studied gene *PRDM9*, which is vital for shaping the recombination landscape (Parvanov et al. 2010). Both introgressed in event B, most likely from the Arboreal to the Terrestrial clade ancestor (Figure S34).

### Introgression deserts contain genes encoding morphological traits

Regions with strong signals of introgression were significantly depleted for several GO-terms related to anatomy and development (e.g., cell differentiation [GO:0030154], animal organ development [GO:0048513], nervous system development [GO:0007399] and anatomical structure morphogenesis [GO:0009653], Table S6). This suggests that genes related to morphological traits may reduce gene flow, maintaining species integrity. To better understand the processes involved in reproductive isolation, we also specifically investigated the gene content of long genomic regions (≥ 500 kb) devoid of introgression, here termed “introgression deserts” (see Methods). Such regions were likely purged of introgressed variants rapidly after hybridization, and may thus contain genes acting as barriers to gene flow. We found 59-135 such regions across all introgression events, containing 3,354 protein-coding genes (n = 391-995 per event, Table S7), without any significant GO-enrichments. In events A, B and G, the proportion of the X-chromosome contained in introgression deserts was substantially larger than for any autosome, suggesting that the X-chromosome plays an important role in reproductive isolation in these events (Figure S35). In the remaining events the introgression deserts were approximately equally common across the autosomes and the X-chromosome, spanning ca. 0-5 % of the chromosomal length. The majority of genes located in introgression deserts were unique to one event (n = 2,647), and none of the event-specific gene lists were significantly enriched for any GO-terms. However, two genomic regions were identified as introgression deserts in five independent events. These contained ten genes, including *HPS6* which is involved in melanogenesis and has been associated with pelage coloration in the African wild dog *Lycaon pictus* (Chavez et al. 2019). Guenons have distinct face colorations, suggested to be important for species recognition and mate choice (Allen et al. 2014). It is thus likely that genes with functions related to pelage coloration are important for maintaining species identity in sympatry. Among genes repeatedly found in introgression deserts are also several genes involved in reproductive traits. For example, *APPBP2* and *USP32* (events A, B and C) are associated with reproductive seasonality in sheep (Martinez-Royo et al. 2017), a trait that is also highly variable in guenons (Campbell et al. 2007).

## Discussion

In this study we use a comprehensive whole genome dataset from a taxonomically diverse group of primates to elucidate evolutionary processes that may have facilitated this radiation. Our results reveal a complex evolutionary history shaped by multiple ancient hybridization events despite the presence of ecological, morphological and karyotypic differences. In addition, we resolve the disputed guenon phylogeny (Perelman et al. 2011; Moulin et al. 2008; Springer et al. 2012; Tosi et al. 2003; Guschanski et al. 2013) and confirm that extensive ancient gene flow is the primary source of previous phylogenetic uncertainties. We present a well-resolved guenon species tree that provides a foundation for future studies of guenon evolution.

In line with recent studies suggesting that ancestral introgression may spark subsequent radiations through the introduction of novel genetic variation (Han et al. 2017; Grant and Grant 2019; Meier et al. 2017; Svardal et al. 2020), we find that several species-rich guenon lineages are descendants of hybridization events. For example, we detect strong signals of gene flow between the ancestors of the Arboreal and Terrestrial clades, both of which subsequently underwent extensive radiations. However, their non-admixed sister genera, *Miopithecus* and *Allenopithecus*, remained largely monotypic (two and one species, respectively (IUCN 2022)). By contrast, the ancestors of the diverse *mona* and *cephus* species groups showed the most pronounced ancestral gene flow and exchanged up to 20% of their genomes. Similar admixture proportions have been reported as ’hybrid speciation’ in other studies (Zhang et al. 2023), and may have provided genetic fuel for subsequent radiations in these species-rich groups. While circumstantial at this point, it is thus possible that rampant ancient hybridizations explain the remarkable diversity of guenons, at least in part. Focal studies of specific ancestral hybridization events, using population-level data sets, may help elucidate the causal role of ancestral hybridization in guenon speciation.

Our results suggest frequent gene flow across guenon lineages with different chromosome numbers. While karyotypic differences can act as reproductive barriers (Giménez et al. 2013; Garagna et al. 2014; Mackintosh et al. 2022; Augustijnen et al. 2023), gene flow across different karyotypes has been demonstrated in the common shrew *Sorex araneus* (Horn et al. 2012) and rock wallabies *Petrogale* spp. (Potter et al. 2015). Furthermore, cross-karyotypic hybridization may generate novel, intermediate karyotypes (Lukhtanov et al. 2020). Considering their extreme karyotypic diversity and highly reticulate evolution, guenons provide an excellent system to better understand the influence of hybridization on chromosomal re-organization, and, in turn, the role of such events in reproductive isolation and speciation. We found several genes with reported functions in chromosomal segregation that show strong signatures of introgression across lineages with different karyotypes (Figure 3, Table S5). They may be involved in facilitating cross-karyotypic gene flow by promoting pairing and recombination across homologous chromosomes despite karyotypic differences. Particularly intriguing is the ancestral introgression of *PRDM9*. This gene is critical for shaping the recombination landscape in mammals (Parvanov et al. 2010) and has been found to rapidly cause sterility in mouse hybrids (Forejt et al. 2021). Homogenization of *PRDM9* through introgression of the Arboreal ancestral *PRDM9* allele into the Terrestrial clade may have promoted hybrid fertility and thus facilitating gene flow.

Mitochondrial introgression has been found in many study systems, including primates (Bailey and Stevison 2021; Zinner et al. 2018), and is more likely to occur across recently diverged lineages as mito-nuclear incompatibilities accumulate rapidly and can act as early contributors to reproductive isolation (Burton 2022; Tobler et al. 2019). In line with previous studies (Guschanski et al. 2013), we identified multiple cases of ancestral mitochondrial introgressions in guenons, several of which occurred across deeply divergent lineages separated by at least 4-5 million years of independent evolution. Biogeographical analyses suggest that mitochondrial introgression commonly occurs from the resident into the dispersing taxon (Toews and Brelsford 2012; Mastrantonio et al. 2016). In guenons, females tend to be philopatric, whereas males disperse at sexual maturity (Campbell et al. 2007). Mitochondrial introgression is thus likely to happen when males at the front of an expanding population hybridize with resident females of a different species, a process sometimes referred to as ‘nuclear swamping’, (Zinner et al. 2018; Sørensen et al. 2023). While the nuclear genome remains homogenized across the population, mainly through male-mediated gene flow, the introgressed mitochondria can persist at high frequency at the expansion front. If such edge populations with high frequencies of introgressed mitochondria subsequently become isolated and speciation is completed in allopatry, the pattern of differential retention of mitochondria in the presence of equal levels of nuclear gene flow may arise. Such a scenario fits well with current and inferred historical distribution ranges of guenon species with introgressed mitochondria, as the ancient mitochondrial introgression events tend to have happened in lineages that likely diverged from the source population through range expansion leading to secondary contact with other species (Guschanski et al. 2013). It is intriguing to speculate whether introgression of mitochondrial and co-adapted nuclear variation, as we found, e.g., in the *Allochrocebus lhoesti*/*preussi* lineage, may have contributed to reproductive isolation from the ancestral population. A similar idea was raised by (Moran et al. 2022), who reported a co-introgression of mitochondrial and co-adapted nuclear genes in swordtail fish, *Xiphophorus* spp., and proposed reproductive isolation arising from mito-nuclear incompatibility between the resulting lineage and its closest sister species. If this is a general pattern, mitochondrial introgressions, particularly between deeply divergent lineages, might be an important mechanism in the early stages of speciation.

An intriguing question in speciation research is: to what extent is the outcome of introgression predictable? Studies analyzing multiple independent hybridizing populations between the same pairs of taxa report highly similar admixed genome compositions (e.g., swordtail fish, *Xiphophorus* spp. (Langdon et al. 2022), sparrows, *Passer* spp. (Runemark et al. 2018), and ants, *Formica* spp. (Nouhaud et al. 2022). Guenons constitute an excellent model to study the predictability of introgression between different taxa in a speciation continuum, which has not been previously done. In contrast to the aforementioned studies, our results reveal introgression landscapes primarily unique for each admixed lineage. This difference could be due to the level of evolutionary divergence among hybridizing lineages. High repeatability and predictability may be present in closely related populations or species, whereas low repeatability may be the hallmark of independent hybridizations between different, deeply divergent lineages. The weak repeatability of the introgression landscapes in guenons suggests that genomic incompatibilities accumulated in different genomic regions and genomic barriers to gene flow are predominantly species-/lineage-specific. It is possible that the karyotypic diversification in guenons has altered their recombination landscape, which is expected to be a critical factor in driving introgression (Edelman et al. 2019; Martin et al. 2019; Langdon et al. 2022). As chromosome-level genome assemblies from non-model organisms become increasingly available, this is an emerging future research topic. Nevertheless, we find an underrepresentation of genes involved in morphological structures and development among introgressed regions (Table S5) and a repeated occurrence of genes involved in pelage coloration and reproductive characteristics among introgression deserts (Table S6), pointing to their functional importance in maintaining species boundaries.

Supporting the notion of repeatability on the level of gene function, we detect an enrichment for immune genes among introgressed loci. Sympatric species are likely to be exposed to similar pathogens. Viruses in particular are known to be a strong selective force in guenons (Svardal et al. 2017), and exchanging genes adapted to circulating pathogens may thus be highly beneficial. Guenons are natural hosts to the simian immune deficiency virus (SIV), which is closely related to the human immunodeficiency virus (HIV). Supporting the importance of adaptively introgressed loci with immune function, we detect *CCR2* in multiple introgression events (B, C and D), a gene involved in SIV response in macaques (Bissa et al. 2023).

We also find that elevated heterozygosity in introgressed genomic regions, a predicted outcome given a positive correlation between recombination rate and introgression, is not a universal pattern. Although the karyotypic diversification of guenons may have altered their recombination landscape, heterozygosity is still expected to be elevated in introgressed regions in the admixed lineage (but not necessarily globally). However, a recent theoretical study suggested that the relationship between recombination and introgression rates may be absent or negative in the early stages of species divergence and only later become positive (Dagilis and Matute 2022). Two out of four events lacking this relationship occurred between closely related lineages (F and G, diverged < 2 My before hybridization) and may thus not have been diverged enough to develop the positive relationship between recombination and introgression. However, we caution that accurately identifying introgressed haplotype blocks from phylogenomic data is challenging, particularly when hybridization is ancient enough so that these blocks are short and contain few informative sites. There may also be other factors obscuring these inferences, e.g., large proportions of ILS.

In summary our study untangles a complex evolutionary history of guenons shaped by extensive ancient introgression. We found gene flow across deeply divergent lineages, suggesting that reproductive isolation in this primate clade may remain incomplete for extended periods despite ecological, morphological and karyotype differences. We note that while hybridization frequently resulted in interspecific transfers of mitochondrial genomes, not a single case of introgression of the paternally inherited Y-chromosome was identified in our dataset. This strongly suggests that male hybrids experienced lower fitness, in line with Haldane’s rule (Haldane 1922), which predicts that post-zygotic barriers will likely be stronger in the heterogametic sex. Our study offers insights into the role and prevalence of ancient gene flow in mammalian radiations and provides an important framework for future studies of this diverse primate group.

## Methods

### Mapping and variant calling

Our dataset consisted of short-read whole-genome sequencing data from 37 individuals (22 guenon species and 2 outgroup species) from different sources (Ayoola et al. 2020; van der Valk et al. 2020; Kuderna et al. 2023); see Table S1 for details). We followed the Genome Analysis Toolkit (GATK) best practices for mapping and variant calling (Van der Auwera et al. 2013). Briefly, we marked adapters with MarkIlluminaAdapters (Picard v. 2.23.4, http://broadinstitute.github.io/picard) and mapped reads to the rhesus macaque (*Macaca mulatta*) reference genome (Mmul_10, GenBank: GCA_014858485.1) using bwa-mem (v 0.7.17). Duplicate reads were marked using MarkDuplicates (v. 2.23.4), and we used HaplotypeCaller (v. 4.1.4.1) and GenotypeGVCFs (v. 4.1.1.0) to call genotypes, including invariable sites. Next, we excluded insertions and deletions (indels) and filtered the remaining calls using VariantFiltration (v. 4.1.1.0) with the recommended exclusion criteria (QD < 2.0, QUAL < 30, SOR > 3, FS > 60, MQ < 40, MQRankSum < -12.5, ReadPosRankSum < - 8.0). Next, we excluded multiallelic sites and repetitive regions annotated in the repeat-masker track for the Mmul_10 reference with bcftools (Li et al. 2009) and set heterozygous genotypes with minor allele support to < 0.25 to no call. Last, we used bcftools to remove sites with coverage below and above one standard deviation (SD) of the genome-wide average or with a fraction of missing genotypes higher than 0.1.

### Phylogenetic Inferences

#### Autosomal and X-chromosomal trees

We used IQTree (v 2.0, (Minh et al. 2020)) to construct gene trees across autosomes and the X-chromosome, using the GTR model with 1,000 rapid bootstraps. We used a window size of 25 kb and sampled one region every 500 kb, as the multi-species coalescence model (see below) assumes independent gene trees without internal recombination. Sites with missing data were filtered out and windows with a filtered length below 10 kb were removed. The filtered gene trees were then used to independently estimate phylogenies for autosomes and the X-chromosome under the multi-species coalescence model with ASTRAL (Zhang et al. 2018). ASTRAL was run on all samples and, after confirming the monophyly of all species, we pruned the tree to include a single sample per species when estimating divergence times. We followed the IUCN taxonomy (IUCN 2022), with the exception that we elevated *C. (mitis) albogularis* to species status, following several recent publications (Shao et al. 2023; Lo Bianco et al. 2017; Detwiler 2019).

We used MCMCTree (Yang 2007) for divergence time estimates. To make the analyses computationally feasible, we generated sequence alignments of 20 randomly sampled genomic windows from the filtered ASTRAL input for the autosomal analyses and ten windows for the X-chromosome. Using a custom python script, genotypes were converted into haploid alignments by selecting a random allele at heterozygous sites. The windows were treated as partitions to allow for rate heterogeneity and we used the approximate likelihood calculation (usedata = 3). We applied fossil data calibrations with uniform prior distributions, using hard minimum and soft maximum bounds (2.5% probability of exceeding upper limit), following (de Vries and Beck 2023), on three nodes: i) the split between *Cercocebus* and *Macaca* (5.33-12.51 MYA), ii) the split between the tribes Papionini (including *Cercocebus* and *Macaca*) and Cercopithecini (6.5-15 MY) and iii) the oldest split within Cercopithecini (6.5-12.51 MYA) (de Vries and Beck 2023). We used the HKY85 substitution model and ran four independent runs of 2,000,000 iterations, sampled every 100, and discarded the first 2,000 samples as burn-in. After confirming that the runs converged at similar age estimates, the four runs were combined and analyzed jointly. Convergence and effective sample size (ESS) were assessed in R (R core team 2017).

#### Mitochondrial genome assembly and mtDNA phylogenic tree

The mitochondrial genomes were assembled using the MitoFinder pipeline (Allio et al. 2020). We trimmed adapter sequences from input reads using Trimmomatic (Bolger et al. 2014), and then ran MitoFinder with the metaspades assembler. Next, assemblies were rotated to the same starting position using a custom python script, and aligned with MAFFT (Katoh et al. 2019). The alignment was visually inspected, and assembly errors from incomplete circularization leading to a short, duplicated sequence were corrected. The cleaned assemblies were then annotated with MitoFinder, using the annotation of *Chlorocebus sabaeus* (NC_008066.1) as a reference, and divided into 42 blocks as follows: Each of the 13 protein-coding genes were divided into 1^st^, 2^nd^ and 3^rd^ codon position resulting in 39 blocks, the tRNAs were concatenated into a single block, and the two rRNAs were kept as two separate blocks. We used PartitionFinder2 (Lanfear et al. 2017) to find the optimal partitioning scheme. First, retaining all samples, we built a maximum likelihood tree with RaxML (Stamatakis 2014), with the GTRGAMMA substitution model and 1,000 rapid bootstraps. The obtained topology was then pruned to keep one sample per species, and divergence times were estimated with MCMCTree, using the likelihood function (--usedata 1). As this function is more computationally intense than the approximate likelihood, we ran four independent runs of 1,000,000 iterations, sampling every 50^th^ and discarded the initial 2,000 samples as burn-in. The four runs converged to highly similar age estimates, and after merging we obtained good ESS (≥ 219) for 20/23 nodes, and acceptable ESS (≥ 181) for all nodes.

#### Y-chromosome phylogenetic tree

We identified males in our study dataset as samples with mapped read coverage on X/Y at approximately half of that across the autosomes. The Y-chromosomal genotypes were then filtered to obtain high-confidence consensus sequences for these samples by excluding sites where any female had a called genotype or any male was heterozygous. The filtered sites were concatenated into a single alignment, and RaxML was run using the same parameters as for the mitochondrial data.

### Gene flow

#### D-statistics

We used Dsuite (Malinsky et al. 2021) to calculate the D-statistics (Patterson et al. 2012) for autosomal bi-allelic SNPs for all possible trios of species in agreement with our inferred species tree (see Results), using the macaque as outgroup. For this analysis we grouped the samples by species and focused only on interspecific gene flow. The only exception was *C. mona*, for which we kept the eastern and western lineages separate, as they were shown to have experienced differential gene flow and mitochondrial introgression with an unknown lineage (Ayoola et al. 2020). A significant D-value indicates gene flow between the mid-group (P3) and either of the in-group sisters P1 (negative D) or P2 (positive D). Since a single hybridization event can cause multiple significant D-values, additional steps were taken to identify hybridization events. First, we used a parsimonious approach to explain all significant D-values with as few gene flow events as possible: if all species of a particular clade produced similar D-values, we interpreted this as gene flow in their common ancestor. Second, we used an approach combining the methods of Dfoil (Pease and Hahn 2015) and the partitioned D-statistic (Eaton and Ree 2013), which incorporates an additional P4 lineage, sister to P3, and offers a more detailed understanding of timing and directionality of gene flow. We used the following rationale: If gene flow occurs from P3 into P2, the shared ancestry between P3 and P4 will lead to significant D-values also between P4 and P2, albeit lower than between P3 and P2. However, if gene flow occurred in the opposite direction, from P2 into P3, there would be no excess allele sharing between P2 and P4. Dfoil uses four different D-statistics (*D_FO_, D_IL_, D_FI_, D_OL_*), where the directionality of gene flow will result in various combinations of positive, negative or non-significant D-values (Pease and Hahn 2015). The partitioned D-statistics (Eaton and Ree 2013) also make use of the shared ancestry between P3 and P4, such that if a positive D-value between P2 and P4 is solely driven by sites introgressing from P3 into P2 (here referred to as ’carryover’ effects), there should be no introgression of alleles emerged on the P4 lineage after the split from P3. Based on both these methods, we calculated D-statistics in a two-step fashion for all significant quartets for which a P4 lineage was available. In step 1 the baseline D-statistics were calculated for the four possible combinations (similar to the *D_FO_, D_IL_, D_FI_* and *D_OL_* of dfoil, except that we only considered sites where P3 had the derived allele, following (Svardal et al. 2020)). In step 2 we re-calculated these D-statistics using only SNPs that were differentially fixed either between P1 and P2 (*D_FO_* and *D_FI_*) or between P3 and P4 (*D_IL_* and *D_OL_*), thus removing any signal caused by shared ancestry. For example we can consider a case of bidirectional gene flow between P2 and P3 (Figure 2). In addition to the positive D-statistics between these lineages (*D_FO_ and D_IL_*) that are the result of direct gene flow, P2 will introduce some alleles into P3 that it shares with P1 returning a positive *D_FI_*, and P3 will, in turn, transfer some alleles to P2 that it shares with P4 (positive *D_OL_*). The positive D-values resulting from these indirect effects disappear after step 2, as the shared alleles are removed.

#### Phylogenetic networks

We used PhyloNetworks (Solís-Lemus et al. 2017), a model-based approach, to independently test for the gene flow events inferred from D-statistics. We used IQTree to construct ML trees in 25 kb windows, separated by 75 kb, to reduce linkage between windows. We applied the GTR model of evolution and 1,000 rapid bootstraps. Windows with less than 5,000 called sites across all samples were discarded. As network approaches quickly become computationally unfeasible with increasing taxa, and only first-level networks can be inferred with this approach (Solís-Lemus and Ané 2016), this method does not allow for testing all the identified events (see results) in a single run. Therefore, we ran PhyloNetworks on subsections of the species tree (subsets of species), both to give an overview of the most pronounced gene flow events and test the support for each (Figures S13-S20). Ten runs were performed with hmax 0-4 to identify the optimal upper limit of hybridization events. Following this, we ran 30 additional runs with the optimal hmax and selected the network with the best likelihood score with a topology concordant with our inferred species tree.

### Reconstructing ancestral chromosome numbers

To identify which of the detected gene flow events occurred across karyotypes, we used ChromEvol, a Bayesian approach implemented in RevBayes (Höhna et al. 2016), to reconstruct the ancestral karyotypic states along the guenon species tree. All lineages/species with known chromosome numbers were used in this analysis (Lo Bianco et al. 2017; Sineo et al. 1986). First, we used equal rates of chromosomal gains and losses (delta ∼ dnExponential(10)) and ran 100,000 iterations, discarding 25 % as burn-in. Second, since the karyotypic diversity of guenons has been attributed mainly to fissions (Moulin et al. 2008), we also performed two runs with higher probability of fissions than fusions (delta ∼ dnExponential(11 and 12)).

### Characterizing and comparing the introgression landscapes

To map the introgression landscape for all inferred gene flow events, we conducted a number of statistical analyses in non-overlapping windows along the reference genome. We chose a window size of 25 kb for these analyses, which we considered adequately short given the ancient timing of gene flow but long enough to retain informative sites. We restricted the analyses to windows with at least 1,000 called genotypes, and constructed neighbor-joining trees for each window using PhyML (Guindon et al. 2010). For each introgression event we then used a custom python script to classify the topology of each window into one of five categories: **1)** ’*((P1/P2),P3)’;* “species tree”, **2)** *’((P2/P3),P1);*’ “ancestral introgression”, **3)** ’*((P1/P3),P2)*’; “ILS control”, **4)** ’*((P1/P3_1),(P2/P3_2))*’; “differential introgression” or **5)** “complicated/other“. Note that the latter two trees only occur in cases where P3 or P2 contain more than one sample (either multiple samples per species or multiple species per group in case of ancestral introgression events). The relative abundance of these five topologies and their distribution along the genome was then visualized in R to infer the spatial distribution of introgressed regions.

We also investigated how local genomic ancestry relates to heterozygosity (as a rough proxy for recombination, see Results), by testing if topology category 1, 2, and 3 was associated with different levels of heterozygosity. Window heterozygosity for each sample was calculated using a custom python script, the effect of topology was tested with ANOVA in R and posthoc Tukey’s tests were applied where significant p-values (p < 0.05) were retrieved. P-values from the Tukey’s test were corrected for multiple testing using the Benjamini and Hochberg (BH) method (Benjamini and Hochberg 1995).

Next, we used the python scripts ABBABABAwindows.py and popgenWindows.py (https://github.com/simonhmartin/genomics_general) to calculate F_D_ and nucleotide divergence (D_XY_), respectively, along the reference genome, for each event. F_D_ is related to the D-statistic and estimates discordant allele-sharing patterns in a four-taxon topology, but is more robust in short genomic windows where data is sparse (Martin et al. 2015). To assess whether windows harbored similar levels of introgression among independent gene flow events, we used R to calculate Pearson’s correlations of F_D_ in all pairwise combinations of gene flow events. To investigate gene content of introgressed regions and to reduce the risk of false positives, we required a number of criteria to be fulfilled: F_D_ ≥ 99^th^ percentile of the genome wide F_D_, topology classified as category 2 (“introgressed”), D_XY_ P2-P3 < D_XY_ P2-P1, and, since F_D_ tend to be higher in regions of low absolute divergence (Martin et al. 2015), D_XY_ P1-P3 > 10^th^ percentile of genome-wide divergence. Consecutive windows were merged, and protein coding genes overlapping and contained within 50 kb up and downstream such regions were identified using the Mmul_10 annotation. The lists of genes were analyzed for gene ontology (GO) enrichments using the online tool PANTHER (Ashburner et al. 2000; Mi et al. 2021). We used the human annotation database, included all annotated protein-coding genes in the macaque annotation as a reference list, and restricted the analyses of GO-terms to biological processes.

We also specifically investigated long genomic regions with low levels of introgression, as this indicates that introgressed alleles were purged rapidly after hybridization and suggests a potential role in reproductive isolation. We call such regions ‘introgression deserts’, defined as a region of at least 500 kb (≥ 20 consecutive windows) with F_D_ below the genome-wide average, allowing for a maximum of 5% of windows with missing data. Gene content was assessed and analyzed in the same way as for highly introgressed regions.

### The effect of introgression on species divergence time estimates

To investigate if introgression affected the species divergence time estimates, we re-calculated divergence times using sequence data from twenty 25 kb windows, randomly sampled from three different sets of windows, depending on the levels of local introgression (F_D_) between *cephus* and *mona*: i) low introgression, F_D_ < 25^th^ percentile, ii) random, iii) high introgression, F_D_ > 75^th^ percentile. MCMCTree was used to estimate divergence times with the same settings as for the species tree.

### Co-evolution/introgression of mito-nuclear genes

To investigate if the co-evolution of nuclear genes facilitated mitochondrial introgression, we focused on the mitochondrial introgression event from the *cephus* group ancestor into *A. lhoesti/preussi*. This event stands out in two ways: i) it occurred across branches separated by at least 5 million years (according to the mitochondrial divergence dates closer to 9 million years), and ii) *A. solatus*, the sister of *A. lhoesti*/*preussi*, retained the ancestral mtDNA genome and thus provides an opportunity for comparative analyses. We compiled a list of 199 nuclear-encoded genes with reported mitochondrial function (Bailey and Stevison 2021; Signes and Fernandez-Vizarra 2018), Table S3). Next, we constructed control gene sets by sampling one gene from the remaining protein-coding genes for each N-mt gene, requiring that they have the same gene length +/-10 %. This process was repeated 100 times with replacement, retrieving 100 control gene sets of 199 genes each. We calculated F_D_ and D_XY_ for each gene set with popgenWindows.py and ABBABABAwindows.py (https://github.com/simonhmartin/genomics_general). Fd was calculated for both *A. lhoesti/preussi* and *A. solatus* as P2, using all members of the *cephus* group as P3, *Allen. nigroviridis* as P1 and *M. mulatta* as outgroup. D_XY_ was calculated across three pairs: *A. lhoesti/preussi* vs. *cephus* group, *A. lhoesti/preussi* vs. *A. solatus* and *A solatus* vs. *cephus* group. If N-mt genes co-evolved with the introgressed mtDNA genome, we expect to find a stronger signal of *cephus* group introgression in these genes compared to control genes in *A. lhoesti/preussi*, but no such pattern in *A. solatus*. This would also lead to lower D_XY_ between *A. lhoesti/preussi* and the *cephus* group but increased D_XY_ between *A. lhoesti/preussi* and their sister *A. solatus* in the N-mt genes.

We also counted private *cephus* group SNPs in *A. lhoesti/preussi* in relation to *A. solatus*. This was done to assess fine-scale variation that may have facilitated the mitochondrial introgression. A custom python script was used to identify and count SNPs in two categories: Category 1: *A. lhoesti/preussi* fixed for the same allele as the *cephus* group, whereas *A. solatus* and *Chlorocebus* spp. are fixed for the alternative allele (a nuclear pattern that is concordant with mtDNA genome ancestry), Category 2: *A. solatus* fixed for the *cephus* group allele, *A. lhoesti/preussi* and *Chlorocebus* spp. fixed for the alternative allele (discordant with mtDNA genome ancestry). The number of SNPs in each category was then compared between N-mt and control genes, and the predicted impact of these mutations was estimated using the Ensembl Variant Effect Predictor (McLaren et al. 2016).

## Acknowledgments

We thank Christophe Escudé, Laurianne Cacheux, and Bertrand Bed’Homme at the Muséum National d’Histoire Naturelle, Paris, for providing guenon cell culture samples, and Jean-Pierre Gautier for tissue samples that form the bulk of data used in this project, as well as Mareike Janiak, Tom van der Valk, Simon Martin and Konrad Lohse for bioinformatic support and helpful discussions. The computations were enabled by resources in projects SNIC 2022/6-325 and SNIC 2022/5-561, provided by the Swedish National Infrastructure for Computing (SNIC) at Uppsala University (UPPMAX), partially funded by the Swedish Research Council through grant agreement no. 2018-05973. The project was supported by the Swedish Research Council VR (2020-03398) to KG, Zoologiska Stiftelse grants to AJ, a UKRI NERC Standard grant (NE/T000341/1) to DdV and RMDB.

## Data and code availability

The sequencing data used in this project is available on the European nucleotide archive (https://www.ebi.ac.uk/ena), under accession numbers as listed in Table S1. Custom scripts used for data analyses are available at https://github.com/axeljen/guenon_phylogenomics.

## Author Contributions

Conceptualization: A.J., K.G., Methodology & analyzes: A.J., K.G., F.S, D.d.V., R.B; Sample acquisition: K.G, L.F.K.K., S.K., I.S.C., J.D.K., A.C.K., K.F., J.R., T.M.B., C.R.; Writing—original draft: A.J., K.G.; Writing—review & editing: all authors.

