## Supplementary figures for "Large-scale phylogenomics uncovers a complex evolutionary history and extensive ancestral gene flow in an African primate radiation"

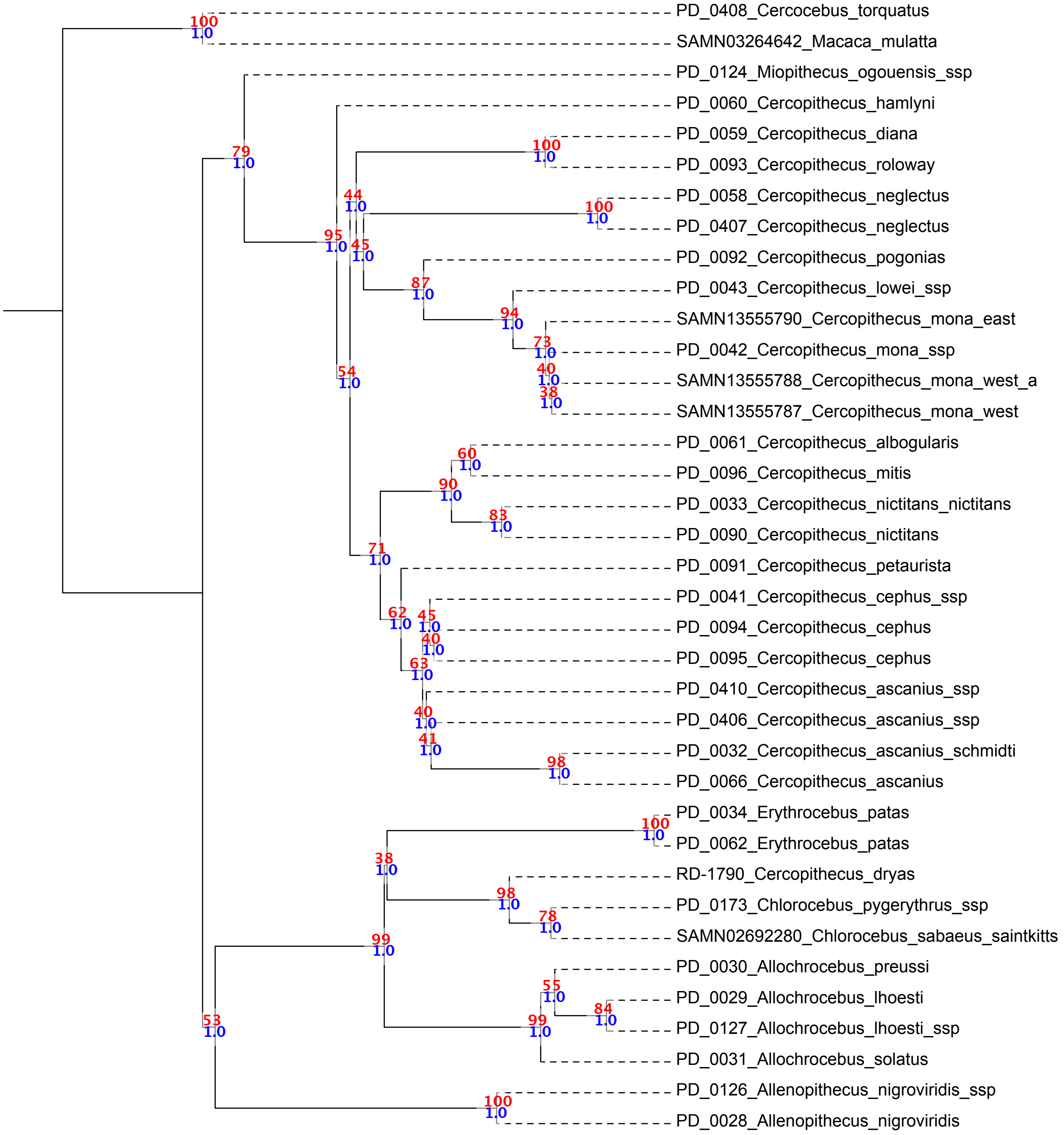


**Fig. S1.**

ASTRAL tree including all samples based on 3,346 autosomal gene trees constructed from one 25 kb genomic window every 500 kb. Blue node annotations show local posterior probability (lpp) support, and red values show the normalized quartet score.

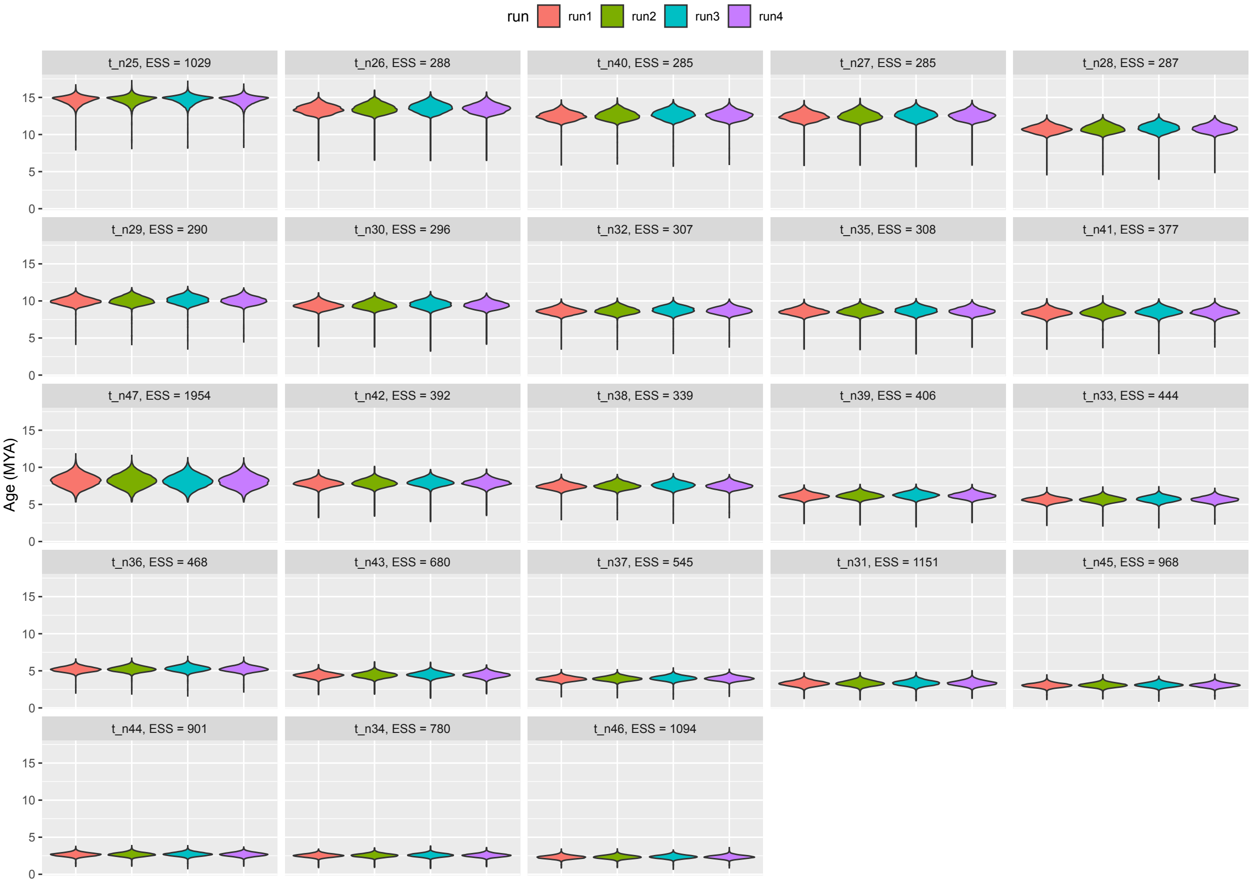


**Fig. S2.**

MCMCTree sample distribution and convergence for autosomal divergence date estimates. Each panel corresponds to one of the 23 internal nodes (as specified in Table S2), and violin plots show the distribution of 20,000 node age samples. The analysis was conducted through four independent runs, shown by the four different colors. Similar age distributions between runs indicate good convergence. Effective sample size (ESS) for the four runs combined is given in each panel header.

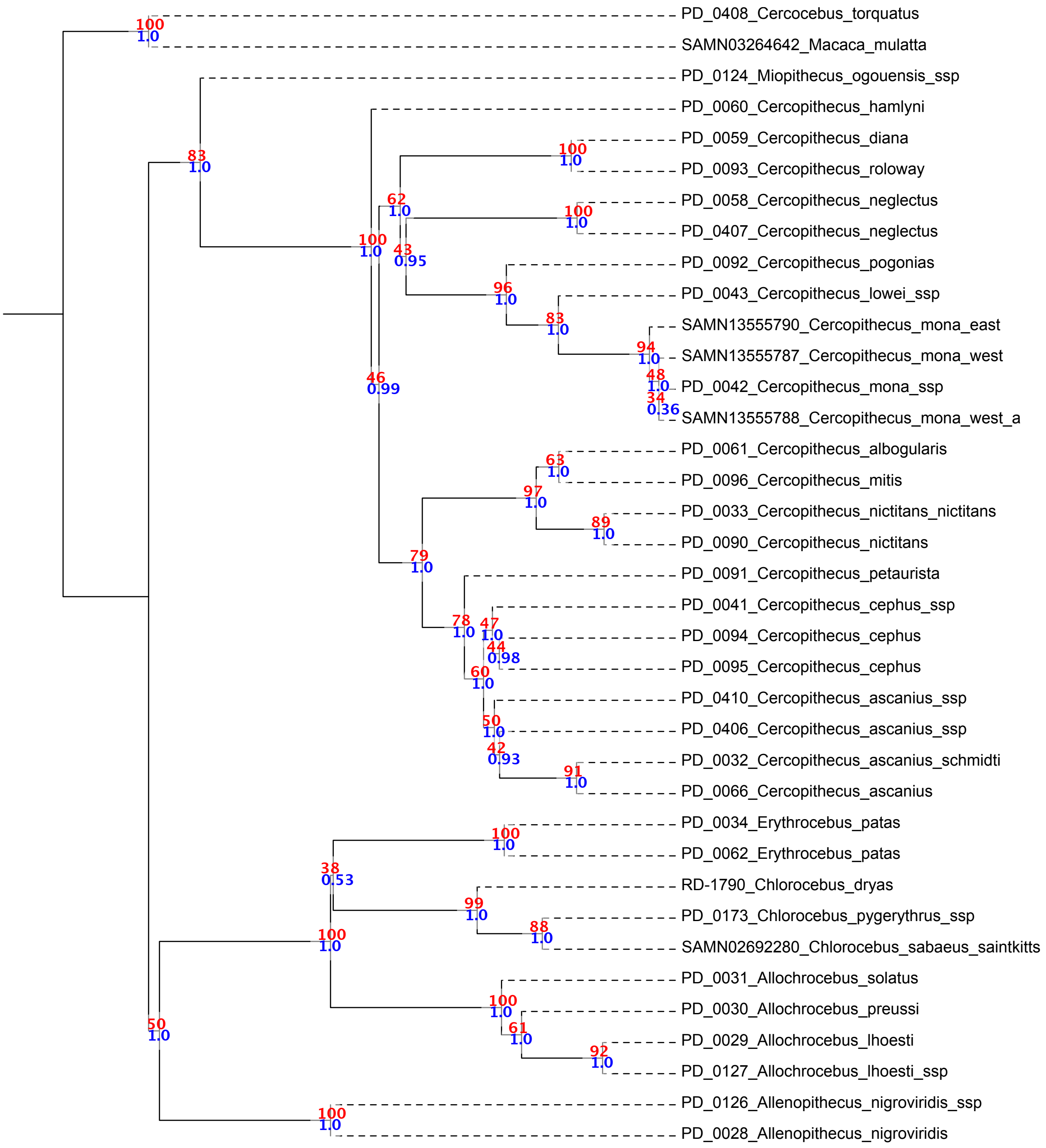


**Fig. S3.**

ASTRAL tree including all samples based on 118 X-chromosomal gene trees constructed from one 25 kb genomic window every 500 kb. Blue node annotations show local posterior probability (lpp) support, and red values show the normalized quartet score.

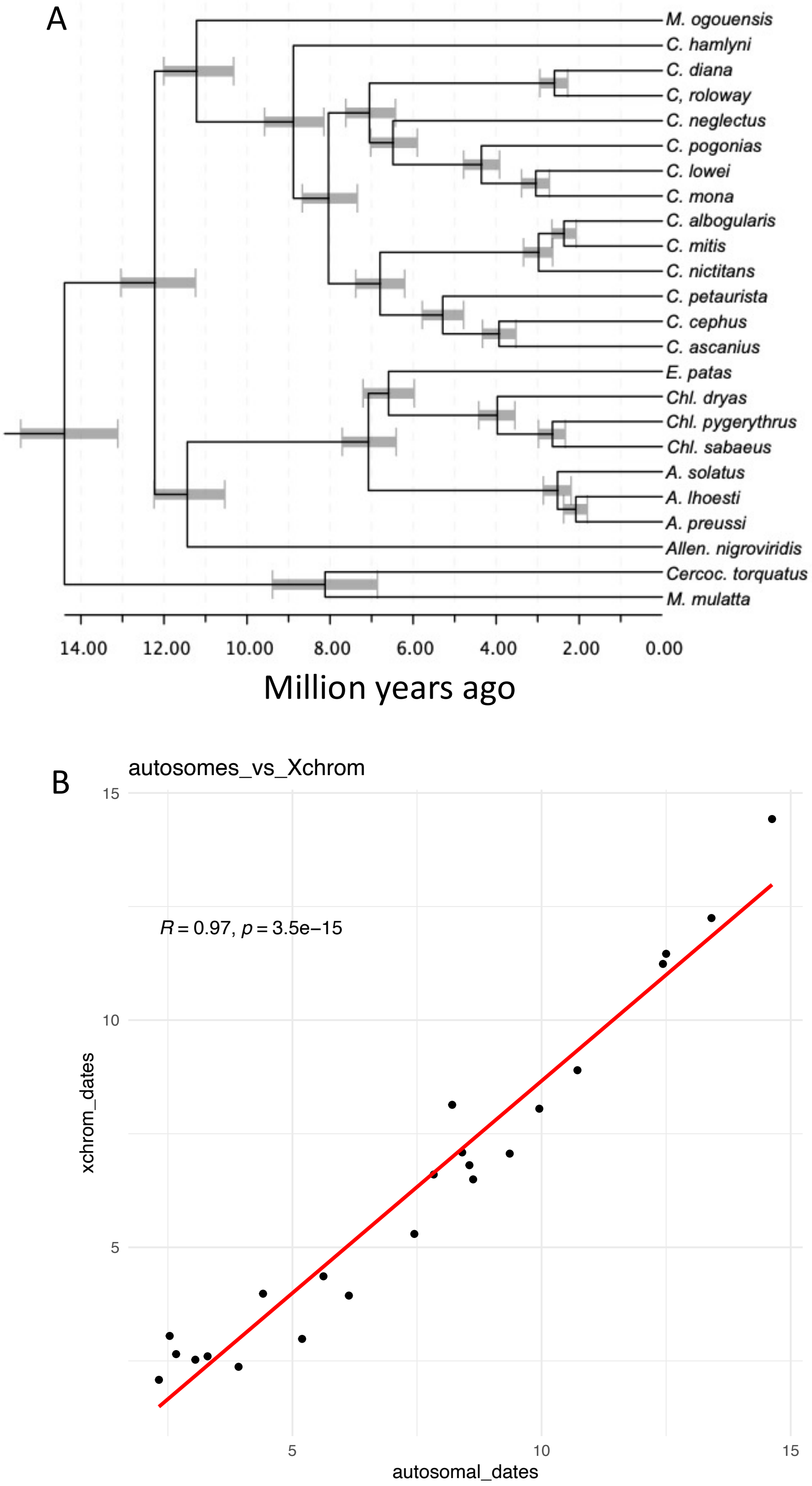


**Fig. S4. (A)** Divergence dates estimated from X-chromosomal loci. **(B)** compares the estimated divergence dates for each node from X-chromosomal data (on the Y-axis) with that obtained from autosomal loci (on the X-axis).

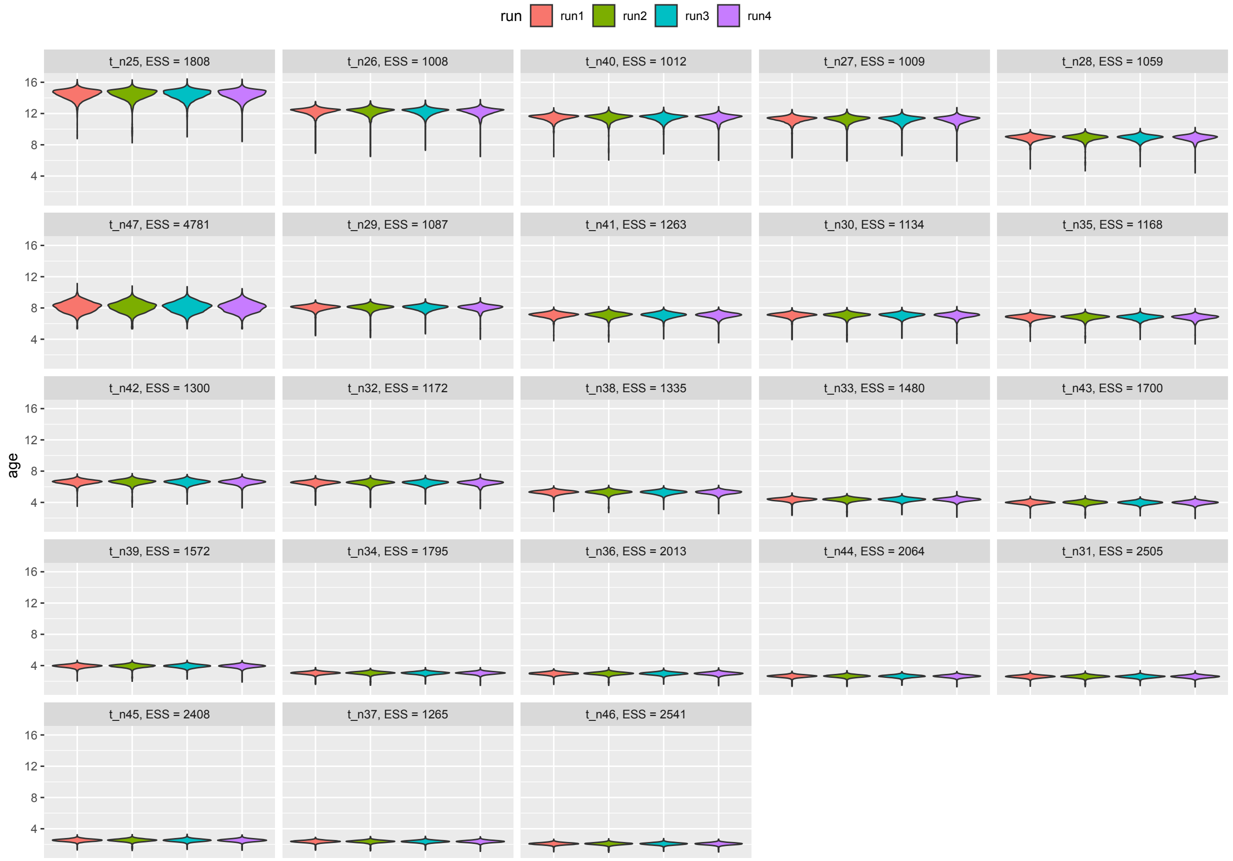


**Fig. S5.** MCMCTree sample distribution and convergence for X-chromosome divergence date estimates. Each panel corresponds to one of the 23 internal nodes (as specified in Table S2), and violin plots show the distribution of 20,000 node age samples. The analysis was conducted through four independent runs, shown by the four different colors. Similar age distributions between runs indicate good convergence. Effective sample size (ESS) for the four runs combined is given in each panel header.

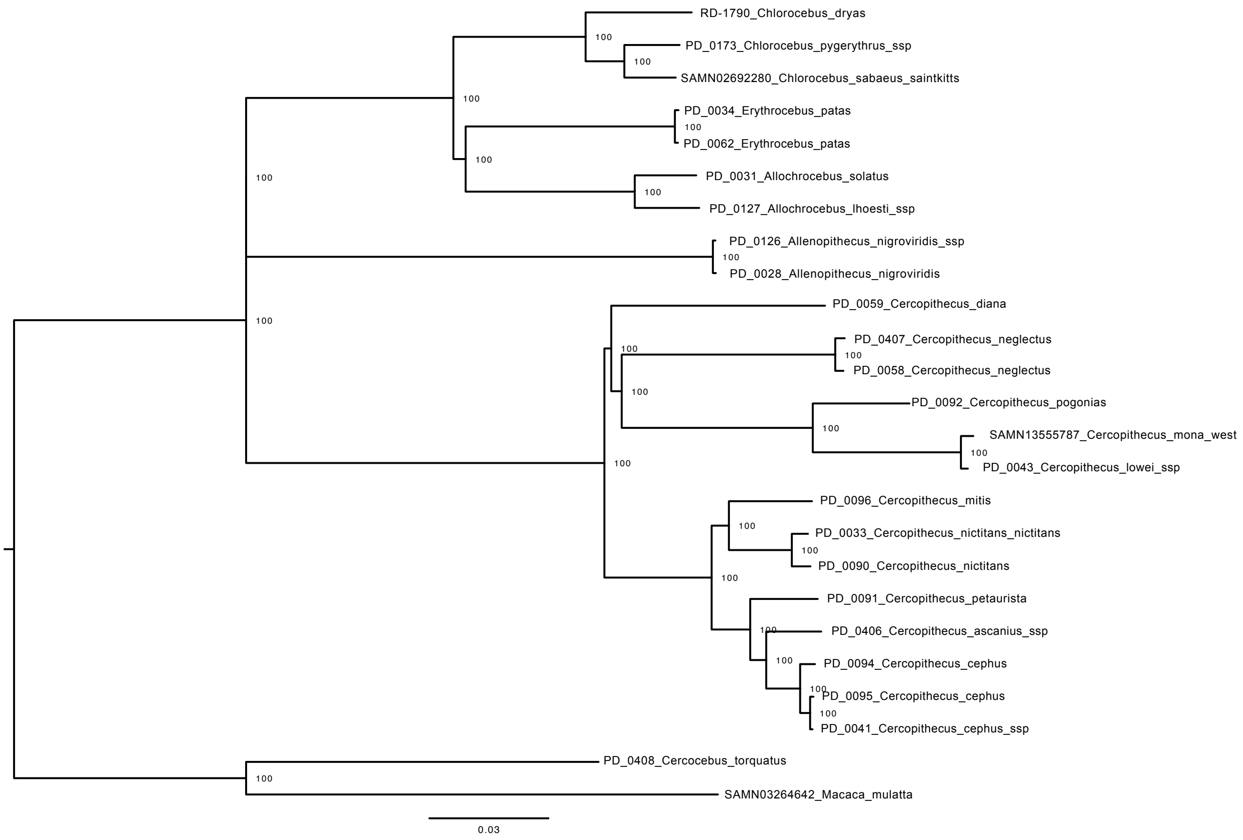


**Figure S6.** RaxML maximum likelihood tree from Y-chromosomal data of all male samples. Node annotations show percentage of bootstrap support (out of 1,000 bootstraps). Topology is identical to the autosomal (main figure 1) with the exeption of *Erythrocebus,* which here is placed as sister to *Allochrocebus*.

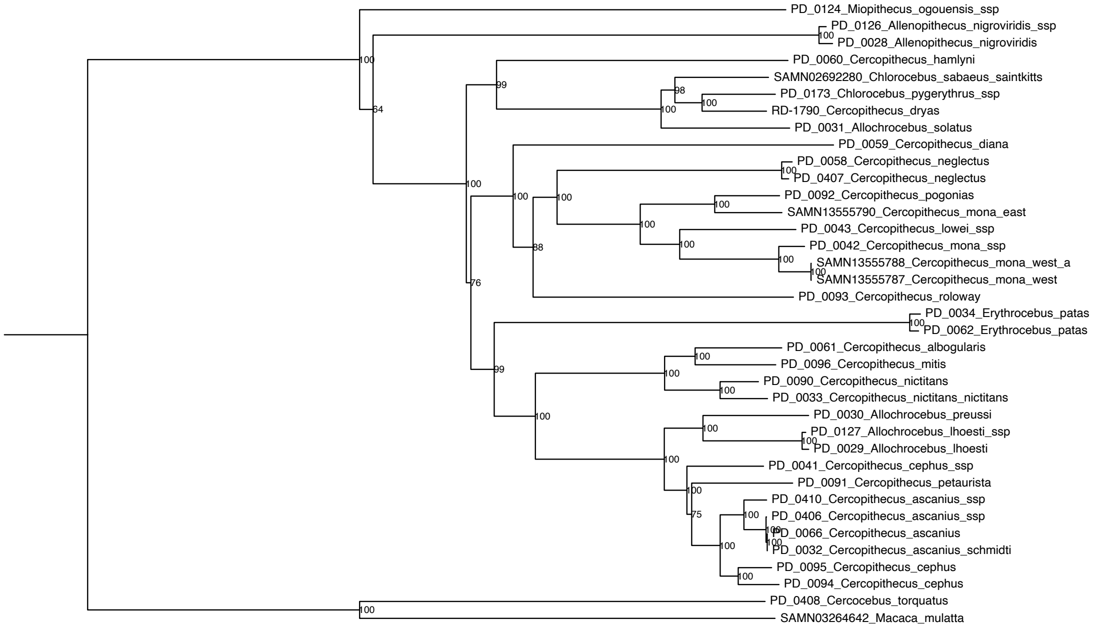


**Figure S7.** RaxML maximum likelihood tree from Y-chromosomal data of all male samples. Node annotations show percentage of bootstrap support (out of 1,000 bootstraps). Topology is identical to the autosomal (main figure 1) with the exeption of *Erythrocebus,* which here is placed as sister to *Allochrocebus*.

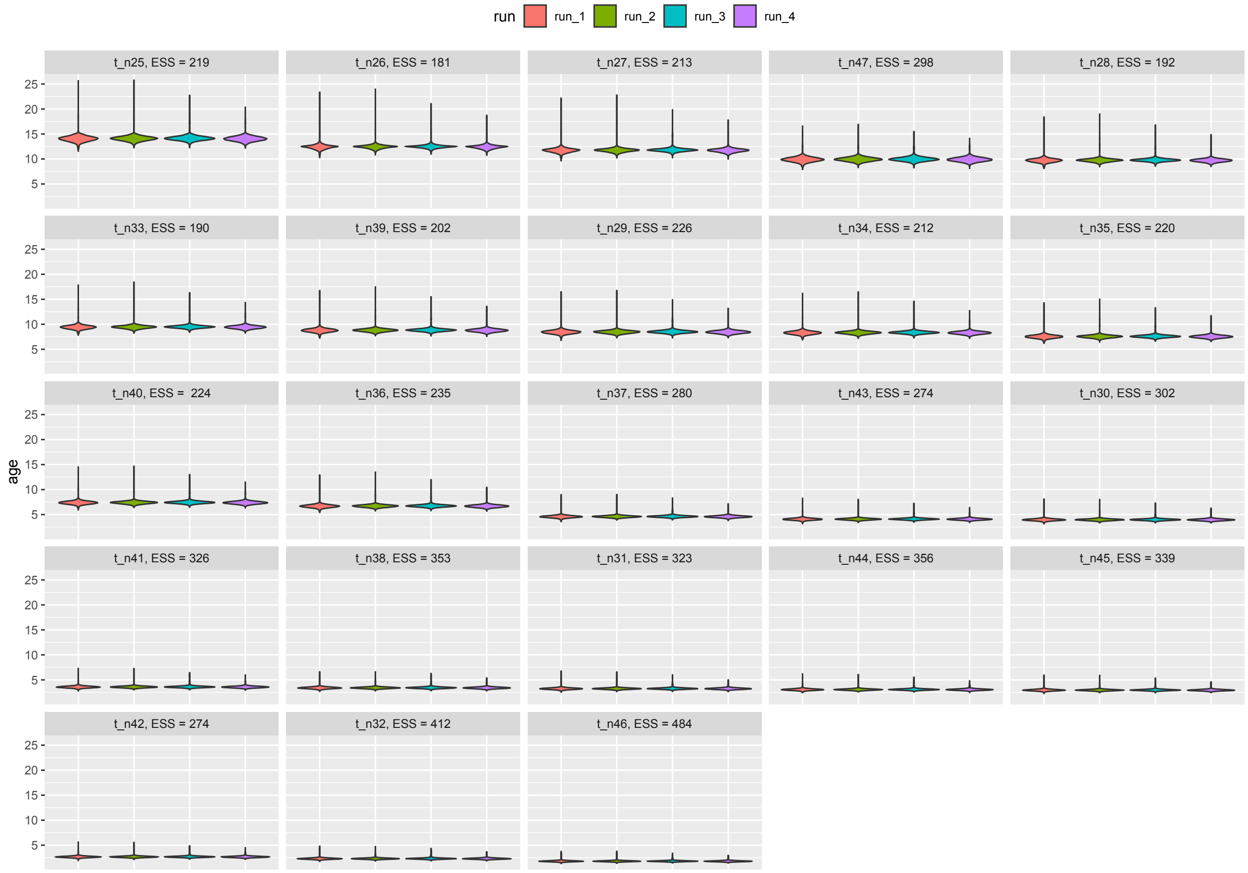


**Figure S8.** MCMCTree sample distribution and convergence for mitochondrial divergence date estimates. Each panel corresponds to one of the 23 internal nodes (as specified in Table S2), and violin plots show the distribution of 20,000 node age samples. The analysis was conducted through four independent runs, shown by the four different colors. Similar age distributions between runs indicate good convergence. Effective sample size (ESS) for the four runs combined is given in each panel header.

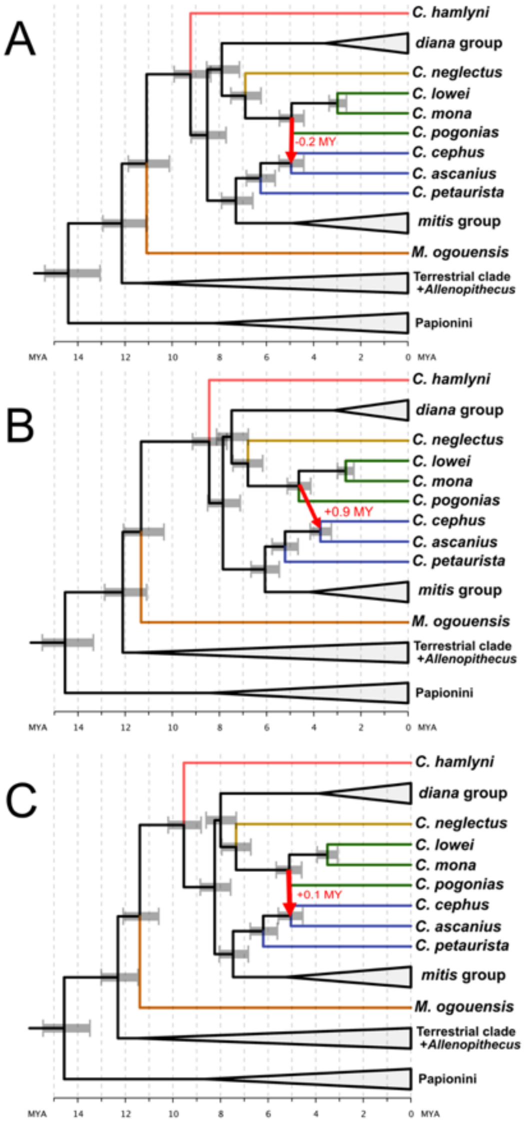


**Figure S9.** Divergence times estimates using 20 randomly selected windows of 25 Kb within three different categories based on introgression level between the *mona* and *cephus* group ancestors, demonstrating the effects of gene flow on divergence time estimates. Panels show divergence dates estimated from 20 windows sampled regardless of introgression level (A), 20 windows sampled from windows with low introgression levels ($F_D < 25^{th}$ percentile) (B), and 20 windows sampled from windows with high introgression levels ($Fd > 75^{th}$ percentile). Panel B provides a considerably younger divergence date between *C. cephus* and *C. ascanius*, making the gene flow from *C. pogonias* into their common ancestor possible.

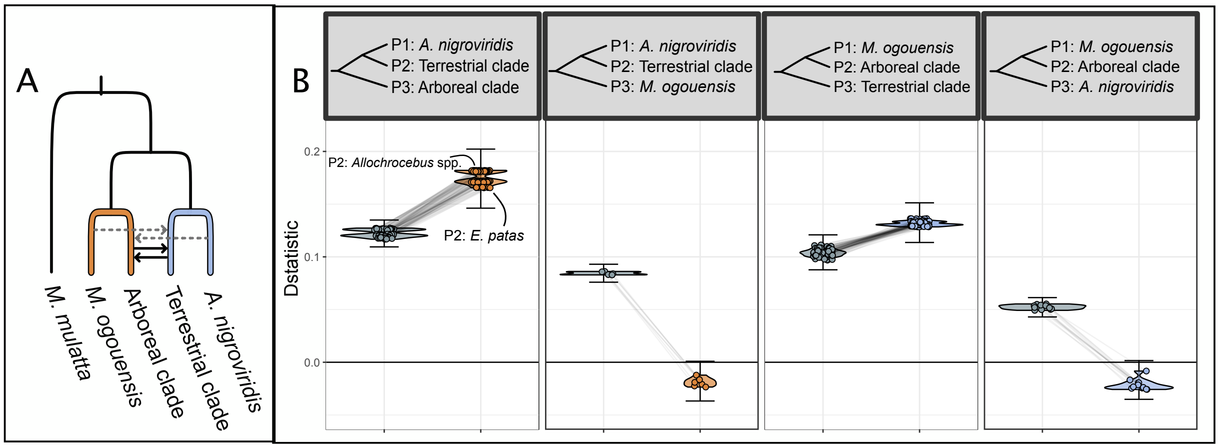


**Figure S10. Excess allele sharing between taxa of the Terrestrial and Arboreal clades.** A) Schematic overview of gene flow event B (main Figure 3), with branch colors corresponding to the partitions used to estimate private allele sharing. Solid black arrows show the inferred gene flow events, and dashed grey lines show 'carry-over' effects. B) Four tests for excess allele sharing between the Terrestrial clade, Arboreal clade, *A. nigroviridis* and *M. ogouensis*. Grey points and distributions show the D-statistics for all combinations of taxa from the respective groups shown in the panel header, with *M. mulatta* as outgroup, using all SNPs. Colored points and violin distributions show the D-statistics for the same trios (connected by grey lines) after removing sites with shared alleles between the Arboreal clade and *M. ogouensis* (orange) or the Terrestrial clade and *A. nigroviridis* (blue). Error bars span the lowest and highest D-statistic +/- 3 standard errors. Differences in the D-statistics after removing shared alleles allow for inferences about directionality: Significant D-statistics driven by shared ancestry are expected to approach zero when shared alleles are removed. Note that some D values turning negative after removing shared ancestral variation is an expected consequence of this method and should not be interpreted as excess allele sharing between P1 and P3 (Pease and Hahn 2015). D-statistics varied somewhat within the Terrestrial clade towards the Arboreal clade, with *Allochrocebus* taxa showing the highest D-statistics, *Chlorocebus* intermediate and *E. patas* the lowest D-statistics, as annotated in the first panel of B.

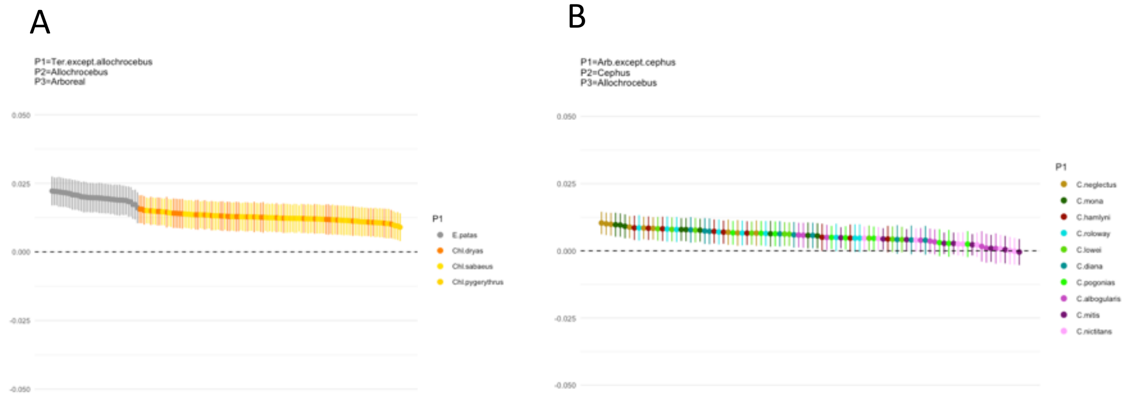


**Figure S11**. (A) Excess allele-sharing between the arboreal taxa and *Allochrocebus* taxa, in relation to the other Terrestrial genera. (B) Excess allele sharing between *Allochrocebus* and specifically the *cephus* group, in relation to other arboreal species.

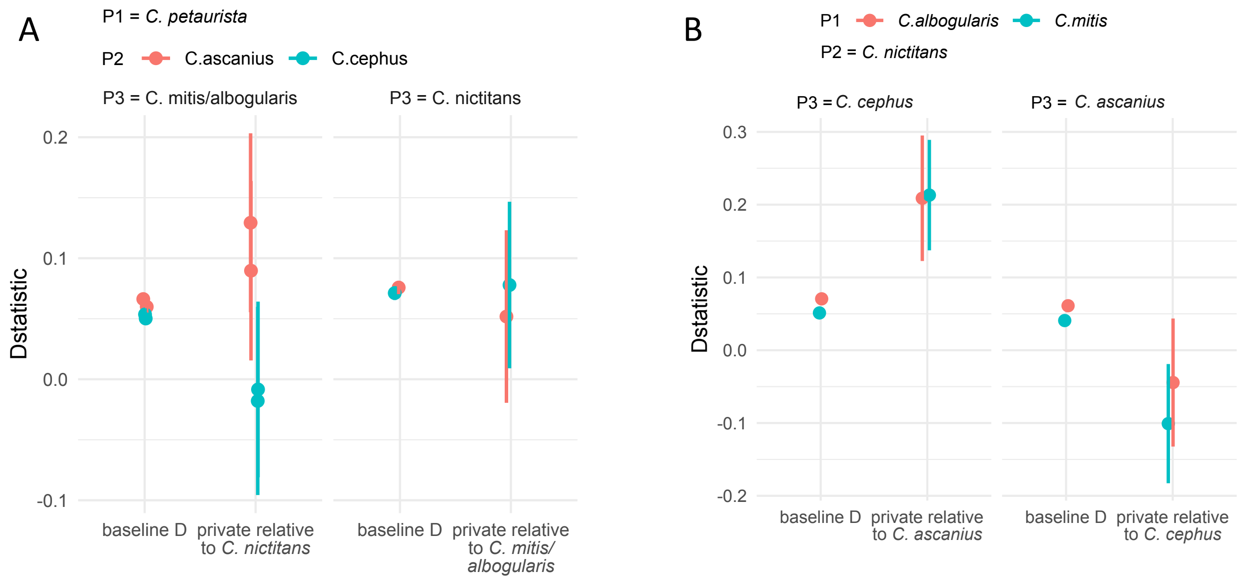


**Figure S12. Excess allele sharing between *C. cephus/ascanius* and the *mitis* species group.** Both baseline (including all SNPs) and private D (shared ancestral variation on the branch leading to different P3 taxa excluded) is shown. A) D-statistics for trios with *C. petaurista* as P1, *C. ascanius/cephus* as P2 and *mitis* group members as P3. *C. mitis/albogularis* shares an excess amount of alleles with both P2 lineages in the baseline D, but only with *C. ascanius* in the private D-statistics. *C. nictitans* shows significant D-statistics with both P2 lineages also in the private analyses, but more so with *C. cephus* than with *C. ascanius*. B) D-statistics for trios with *C. albogularis/mitis* as P1, *C. nictitans* as P2 and *C. cephus/ascanius* as P3. Both *C. cephus* and *ascanius* show significant baseline D-statistics towards *C. nictitans*, but for *C. ascanius* these approach zero in the private D-statistics. These results suggest a complex scenario involving at least two pulses of gene flow: One between *C. mitis/albogularis* ancestor and *C. ascanius,* where directionality can be inferred from the former to the latter due to 'carry-over' effects from *C. cephus* that approach zero when shared alleles are excluded (left facet of panel A). Another pulse of gene flow likely occurred from *C. cephus* into *C. nictitans*.

A

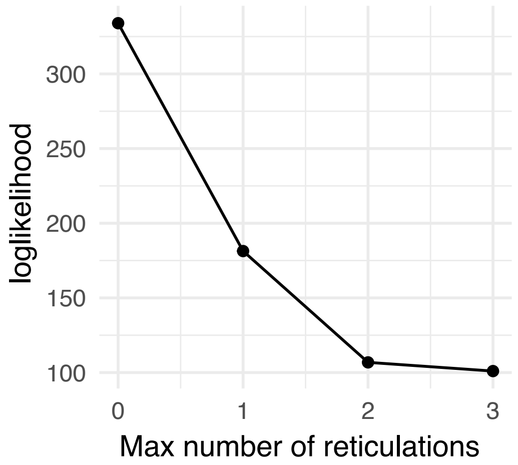


B

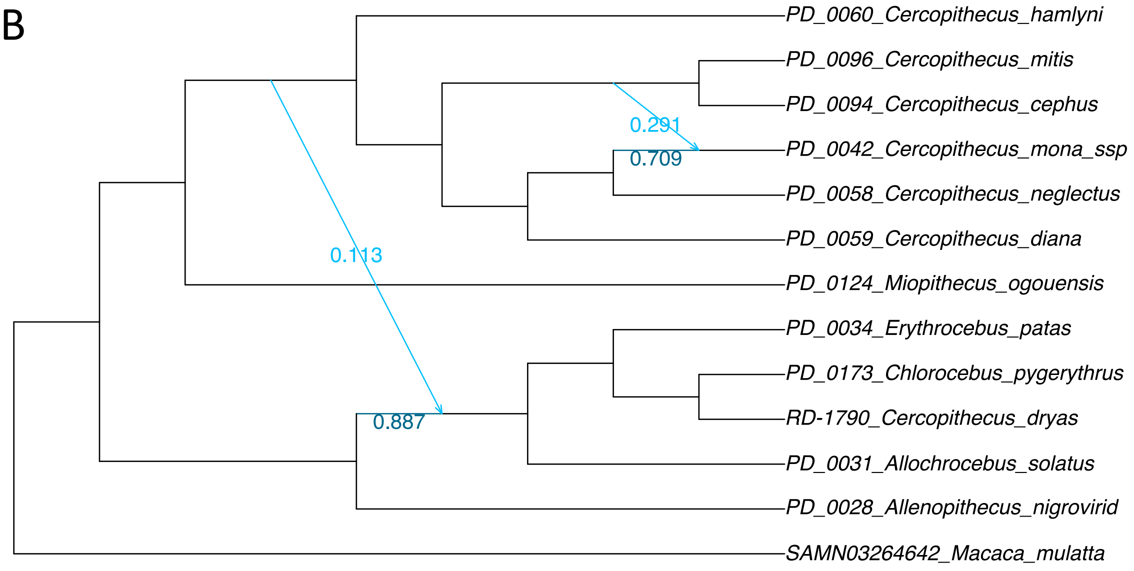


**Figure S13**. PhyloNetworks inferences based on a single representative per species group/genus. A) Loglikelihood for 0-3 hybridization events. B) Outcome of two hybridization events based on (A). Note that the inferred hybridizations here are very similar to the two strongest events identified with D-statistics (events A and B, Figure 3), with the exception that PhyloNetworks places the second hybridization event on the ancestral *mitis*/*cephus* branch, as opposed to the cephus branch in D-statistics.

A

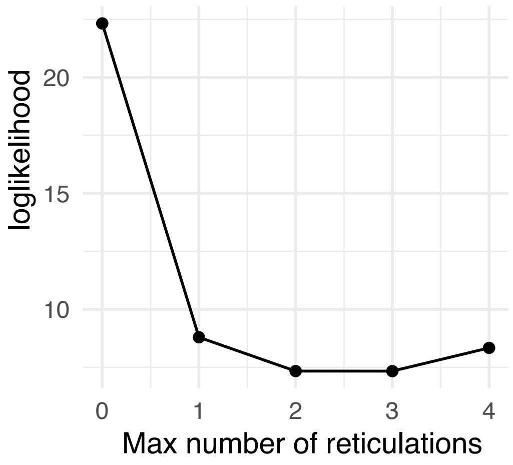


B

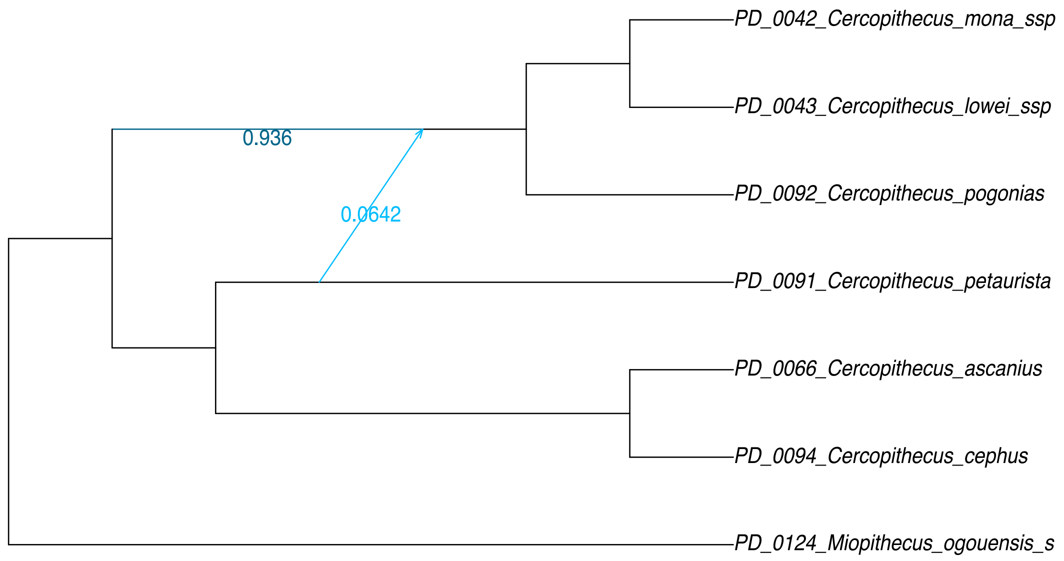


**Figure S14**. PhyloNetworks inferences based on samples from the *mona* and *cephus* species groups, including *Miopitechus* as an outgroup. A) Loglikelihood for 0-4 hybridization events. B) The outcome of one hybridization event based on (A). The inferred hybridization here corresponds to event A1 in the main analysis (Figure 3).

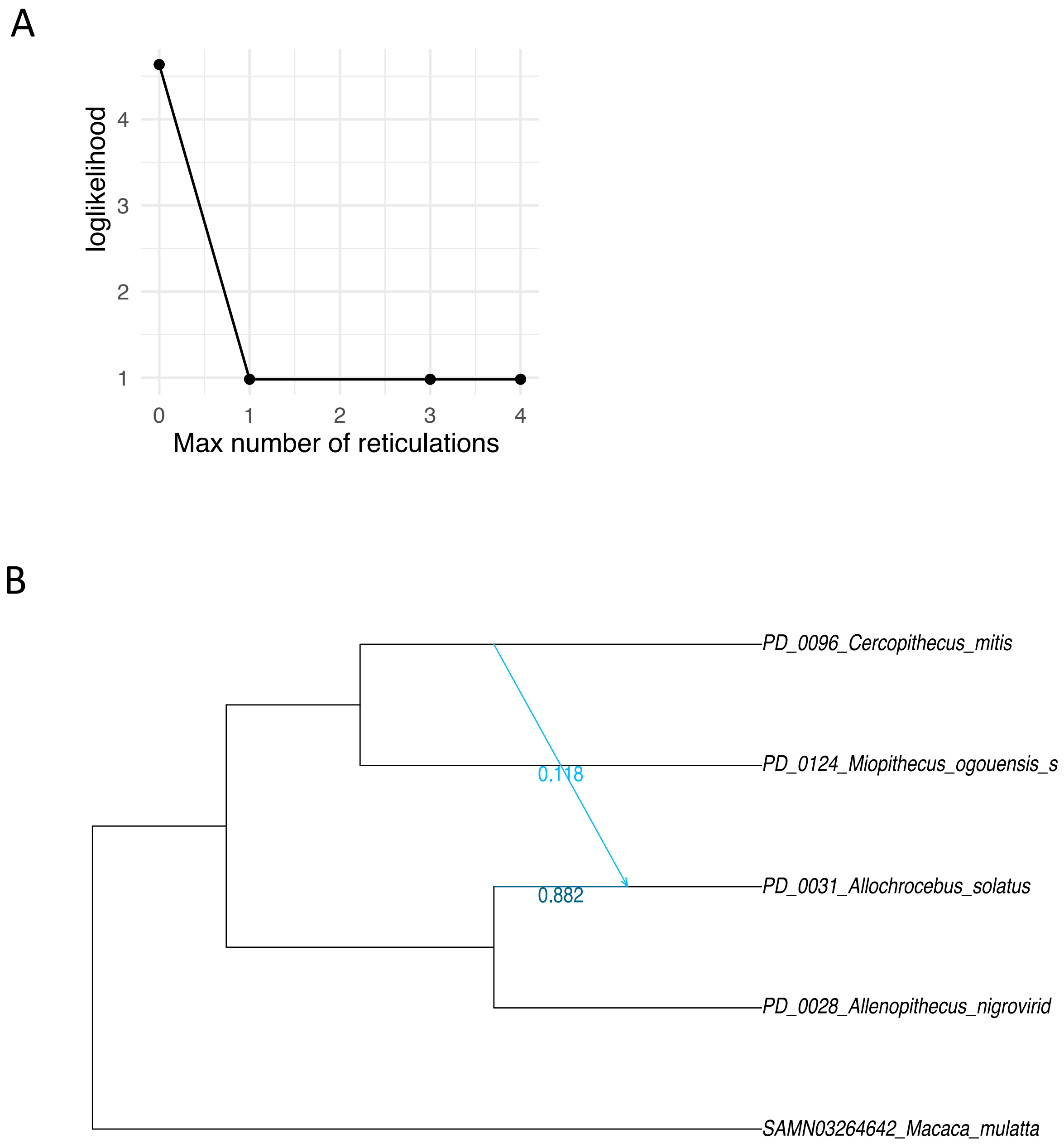


**Figure S15**. PhyloNetworks inferences based on samples from one Arboreal clade lineage (*mitis*), one Terrestrial clade lineage (*Allochrocebus*) and their respective sisters *Miopithecus* and *Allenopithecus*. A) Loglikelihood for 0-4 hybridization events. B) The outcome of one hybridization event based on (A). The inferred hybridization here corresponds to event B in the main analyses (Figure 3, and is also inferred in Figure S9).

A

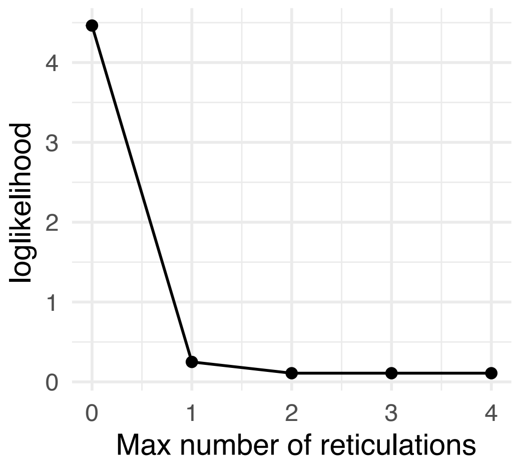


B

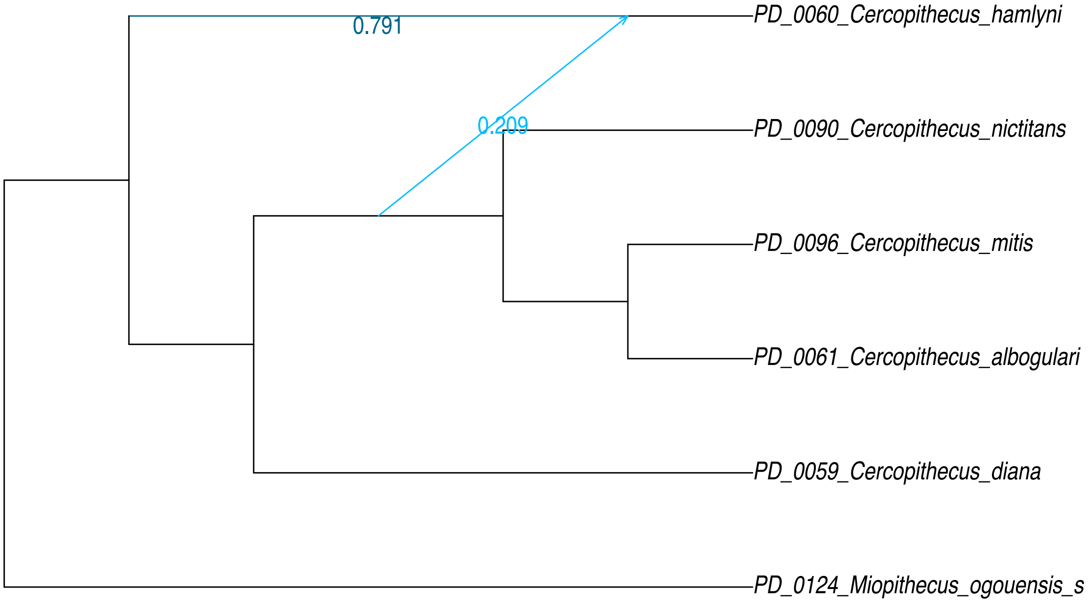


**Figure S16**. PhyloNetworks inferences based on samples from the *hamlyni*, *mitis, diana* groups and *Miopithecus*. A) Loglikelihood for 0-4 hybridization events. B) The outcome of one hybridization event based on (A). The inferred hybridization here corresponds to event C in the main analysis (Figure 3).

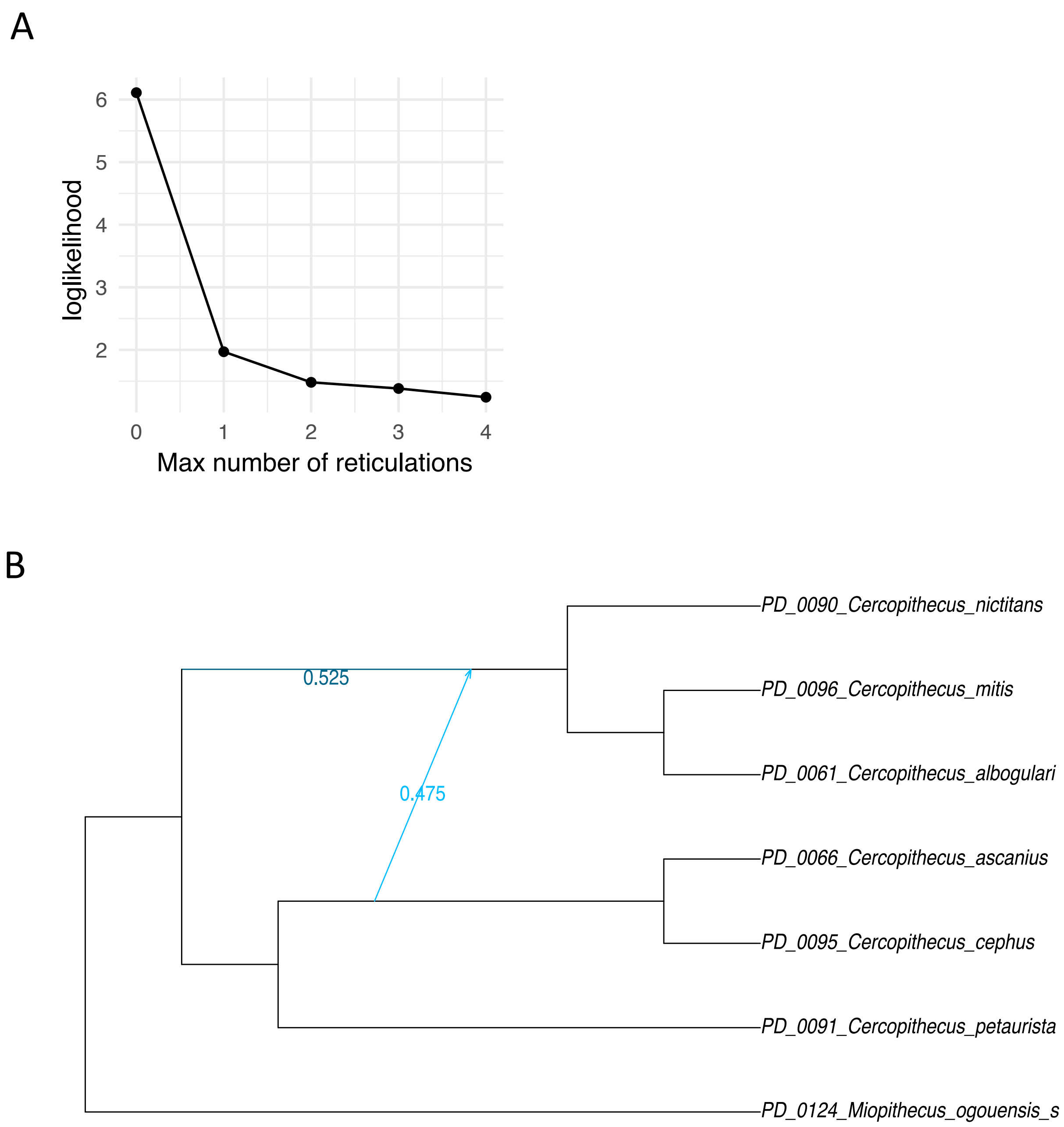


**Figure S17**. PhyloNetworks inferences based on samples from the *mitis* and *cephus* groups and *Miopithecus*. A) Loglikelihood for 0-4 hybridization events. B) The outcome of one hybridization events based on (A). The inferred hybridization here corresponds to event D in the main analysis (Figure 3).

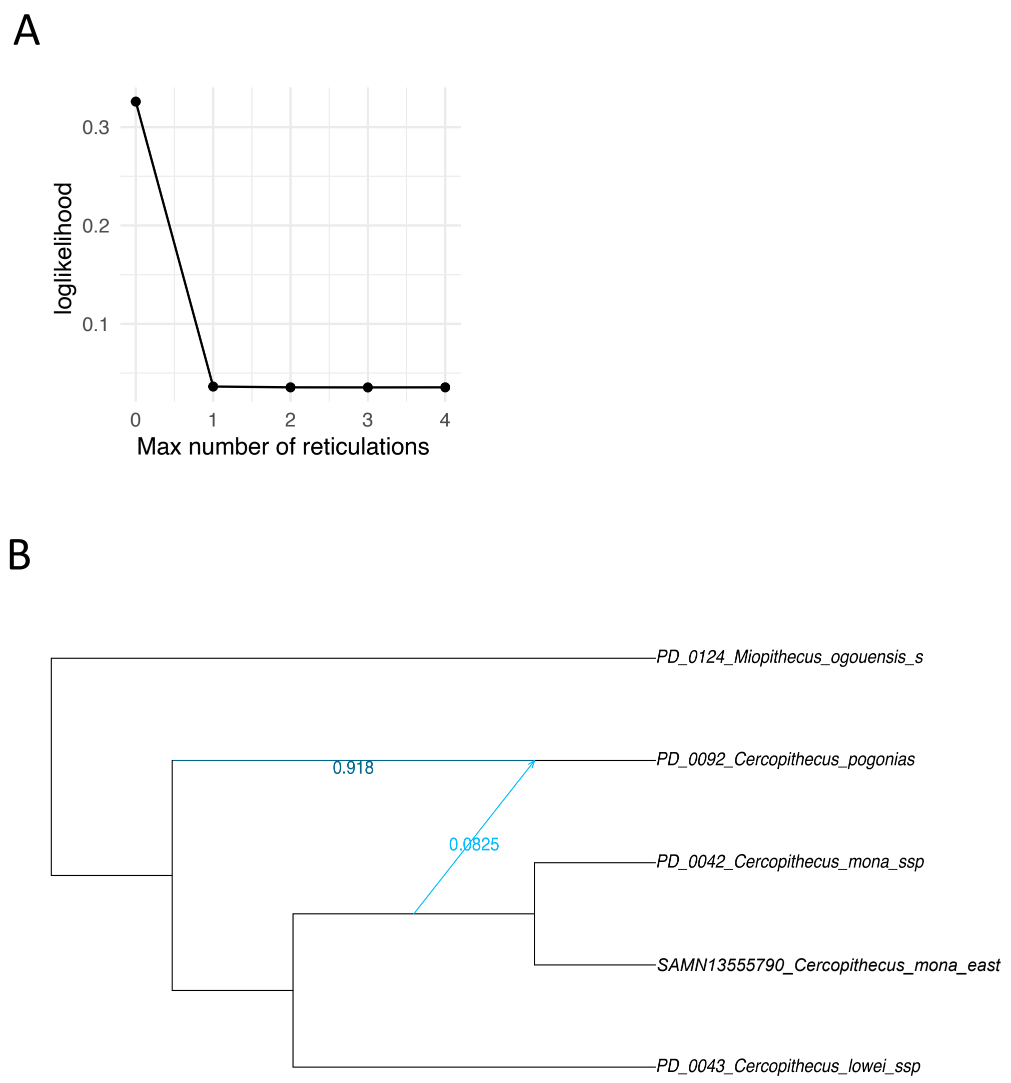


**Figure S18**. PhyloNetworks inferences based on samples from the *mona* group and *Miopithecus*. A) Loglikelihood for 0-4 hybridization events. B) The outcome of one hybridization event based on (A). The inferred hybridization here corresponds to event E in the main analysis (Figure 3). Note that while this arrow is placed on the ancestral *C. mona* branch, our main analyses suggest that it was likely only the eastern population of *C. mona* (here represented by SAMN13555790) that hybridized with *C. pogonias*.

A

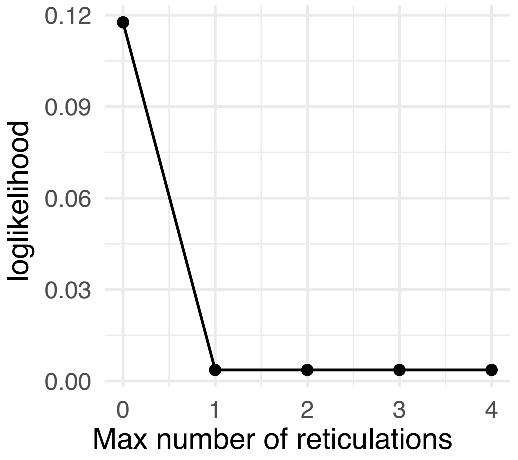


B

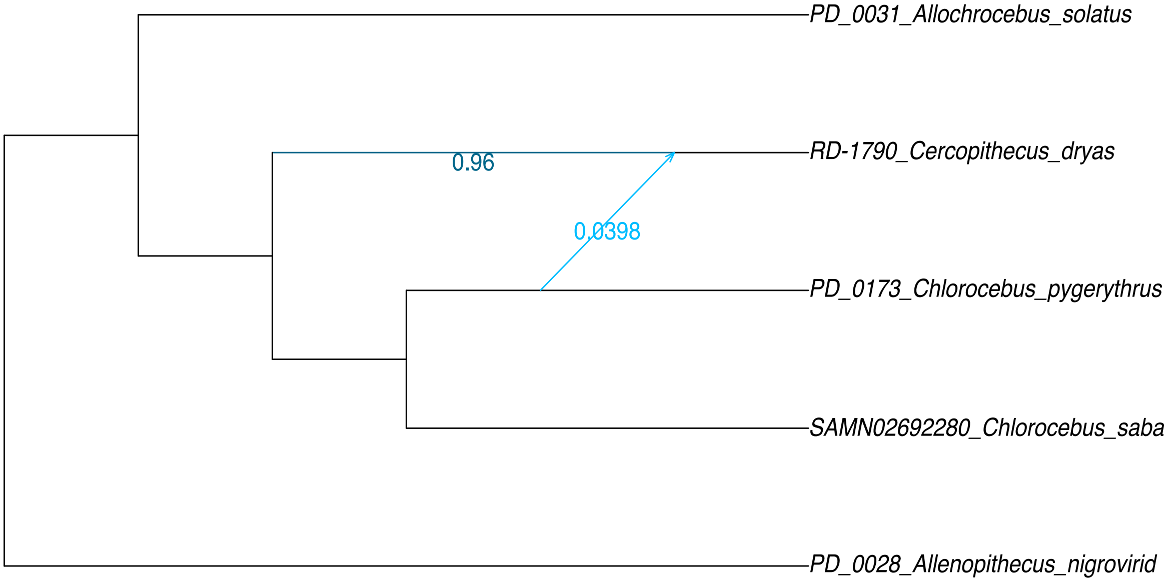


**Figure S19**. PhyloNetworks inferences based on samples from *Allochrocebus, Chlorocebus* and *Allenopithecus*. A) Loglikelihood for 0-4 hybridization events. B) The outcome of one hybridization event based on (A). The inferred hybridization here corresponds to event F in the main analysis (Figure 3).

A

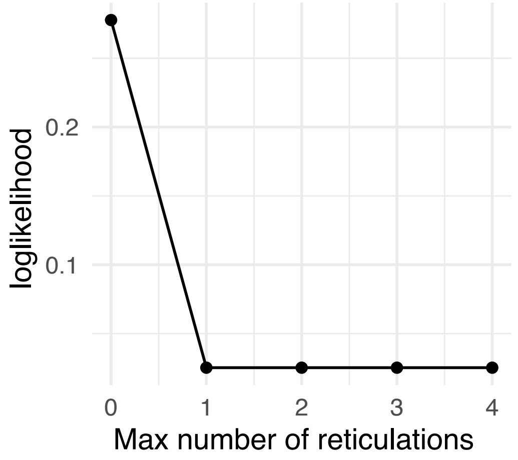


B

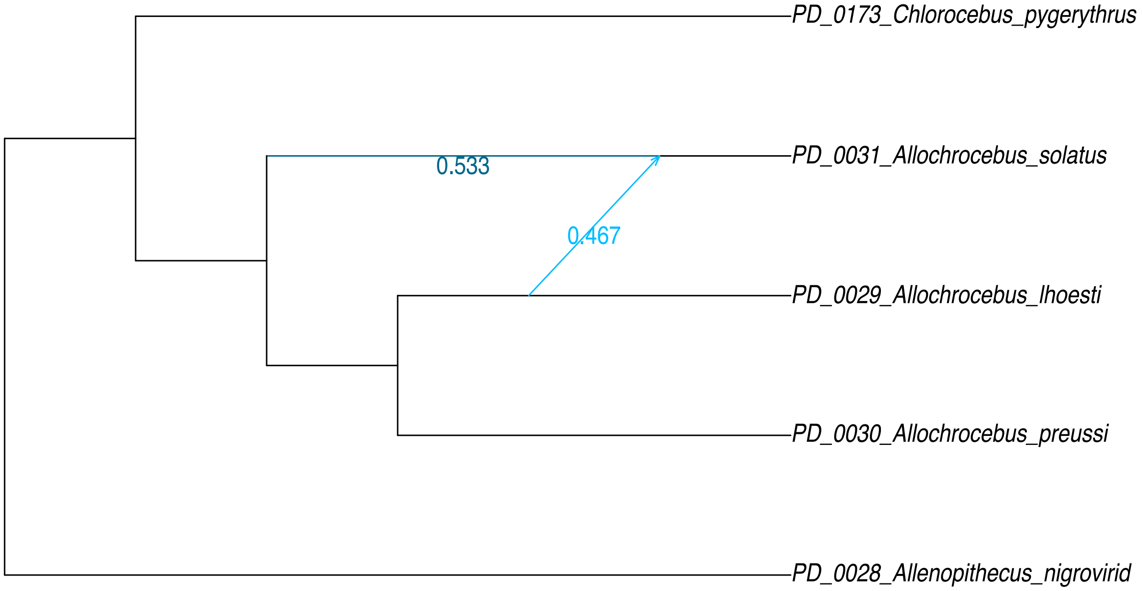


**Figure S20**. PhyloNetworks inferences based on samples from *Allochrocebus, Allenopithecus* and a single sample from *Chlorocebus*. A) Loglikelihood for 0-4 hybridization events. B) The outcome of one hybridization event based on (A). The inferred hybridization here corresponds to event G in the main analysis (Figure 3).

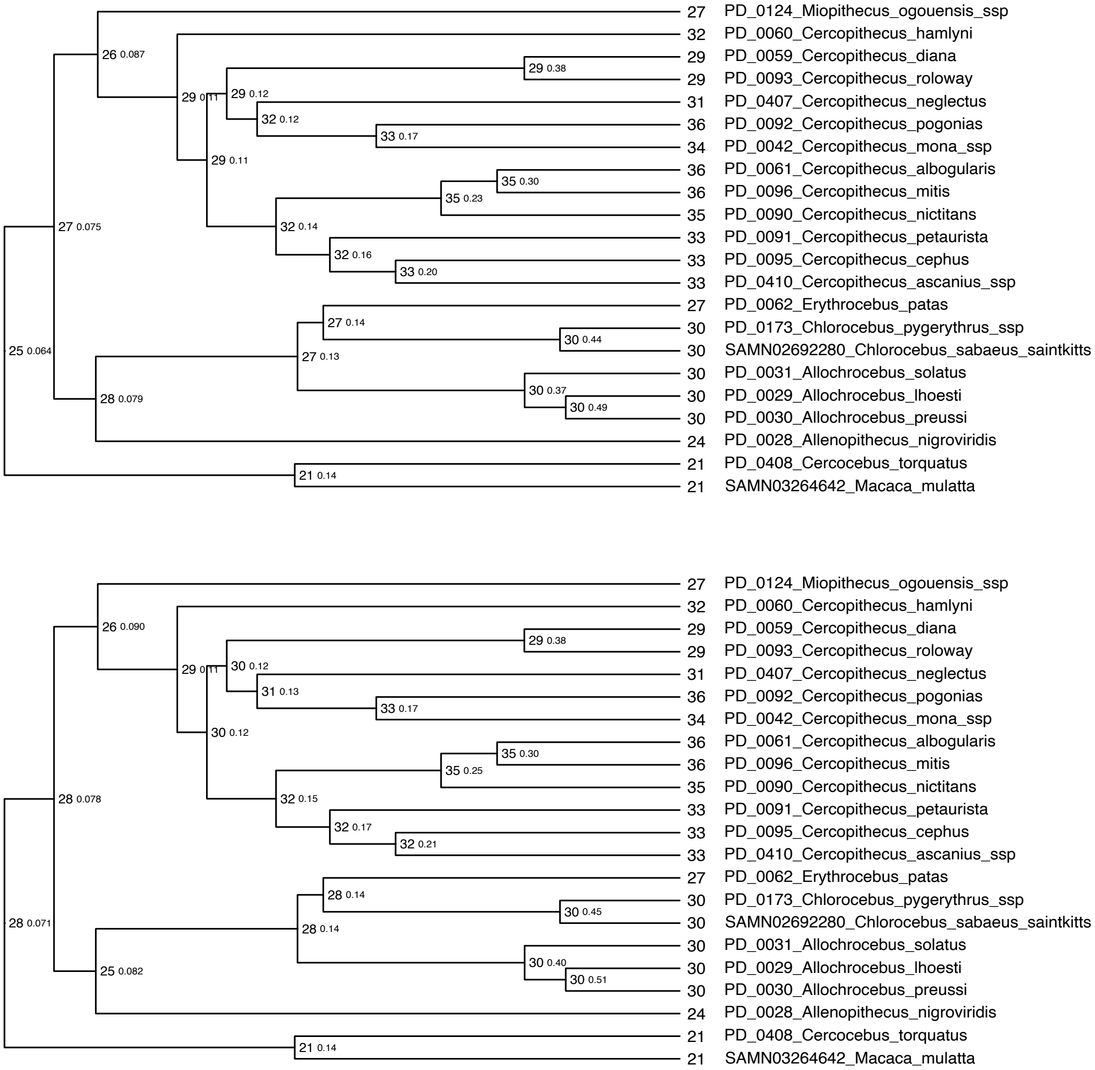


**Figure S21.** Reconstruction of ancestral karyotypes. Large node annotations show the inferred haploid chromosome number, small values show the local posterior probability. Two independent runs of ChromEvol are shown, ran with the same probabilities of fissions and fusions (gamma and delta ~ dnExponential(10.0), default).

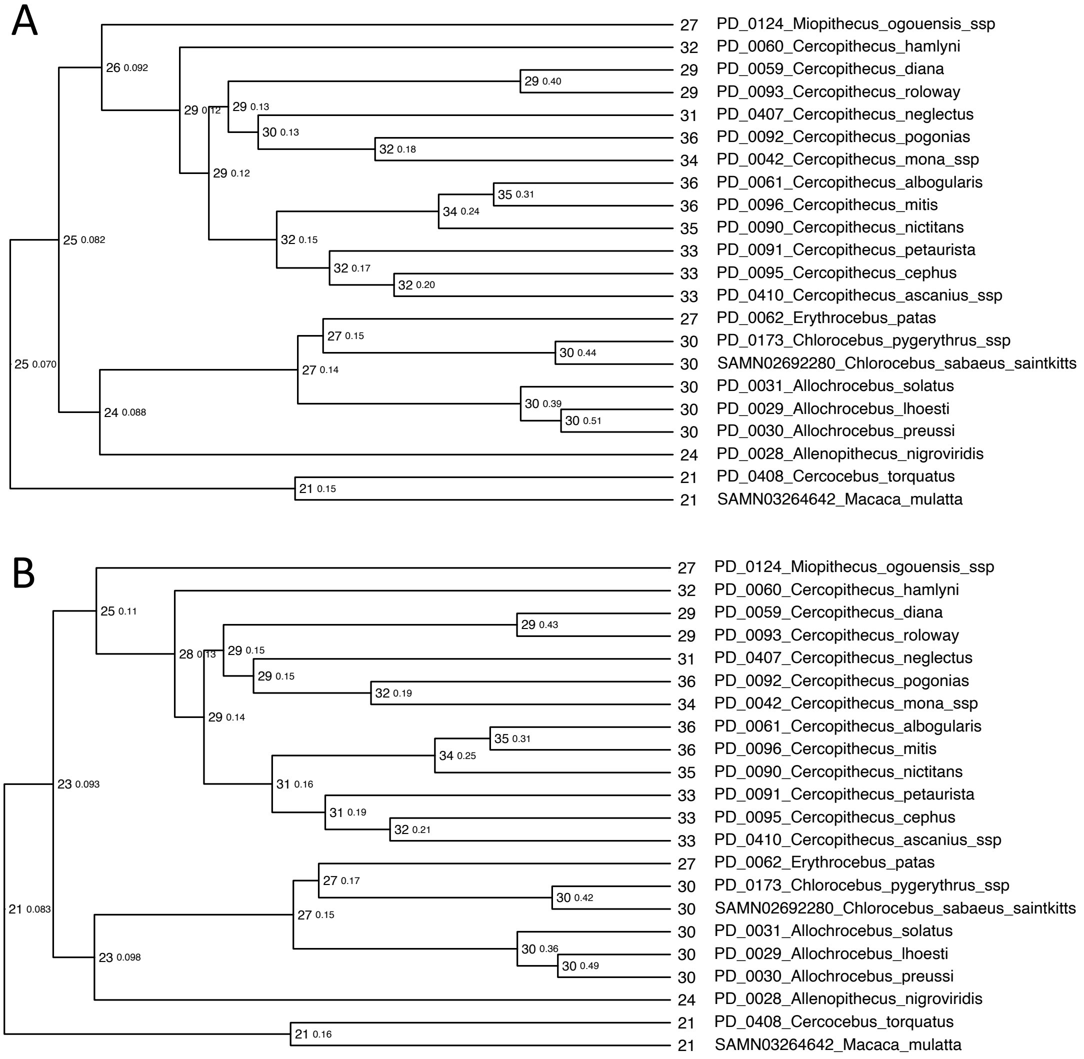


**Figure S22.** Reconstruction of ancestral karyotypes. Large node annotations show the inferred haploid chromosome number, small values show the local posterior probability. Two runs with higher rates of fissions than fusions: gamma ~ dnExponential(10) in both, delta ~ dnExponential(11.0) in (A) and delta ~ dnExponential(12.0) in (B).

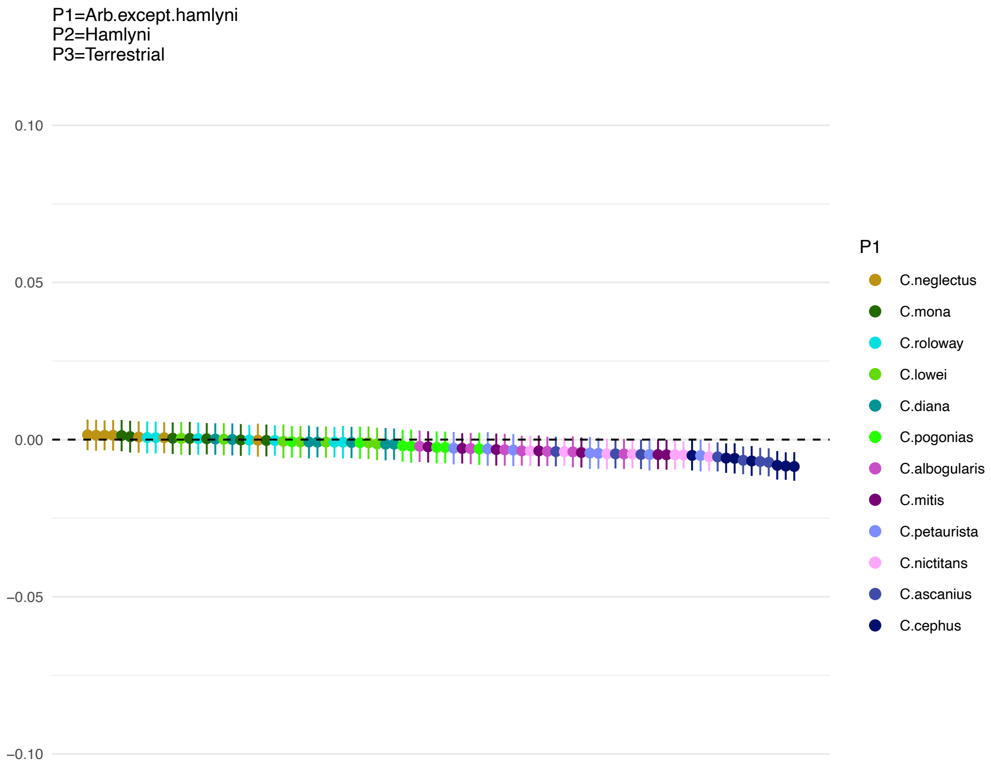


**Figure S23**. ***Cercopithecus hamlyni* did not experience additional gene flow with the Terrestrial clade.** D-statistics between *C. hamlyni* and the Terrestrial clade in relation to other Arboreal clade taxa, showing that there is no evidence for additional gene flow into *C. hamlyni* after the split from the other Arboreal clade taxa.

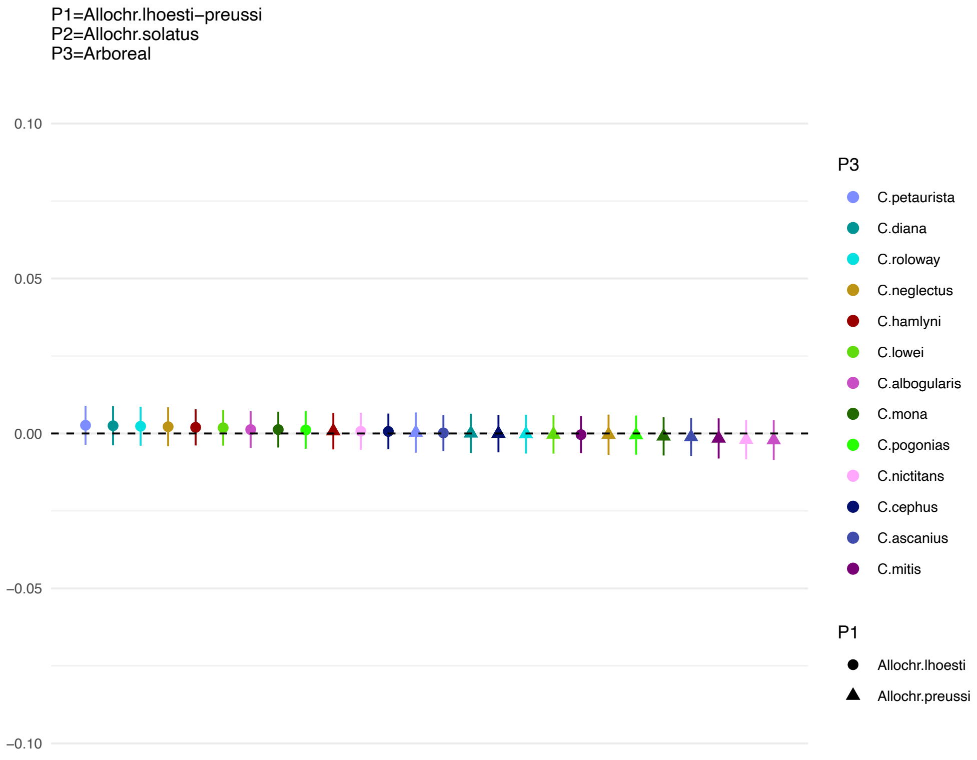


**Figure S24**. D-statistics between *Allochrocebus lhoesti/preussi* and the Arboreal clade taxa in relation to *Allochrocebus solatus*, showing that there is no evidence for additional gene flow into *A. lhoesti/preussi* after they split from *A. solatus*.

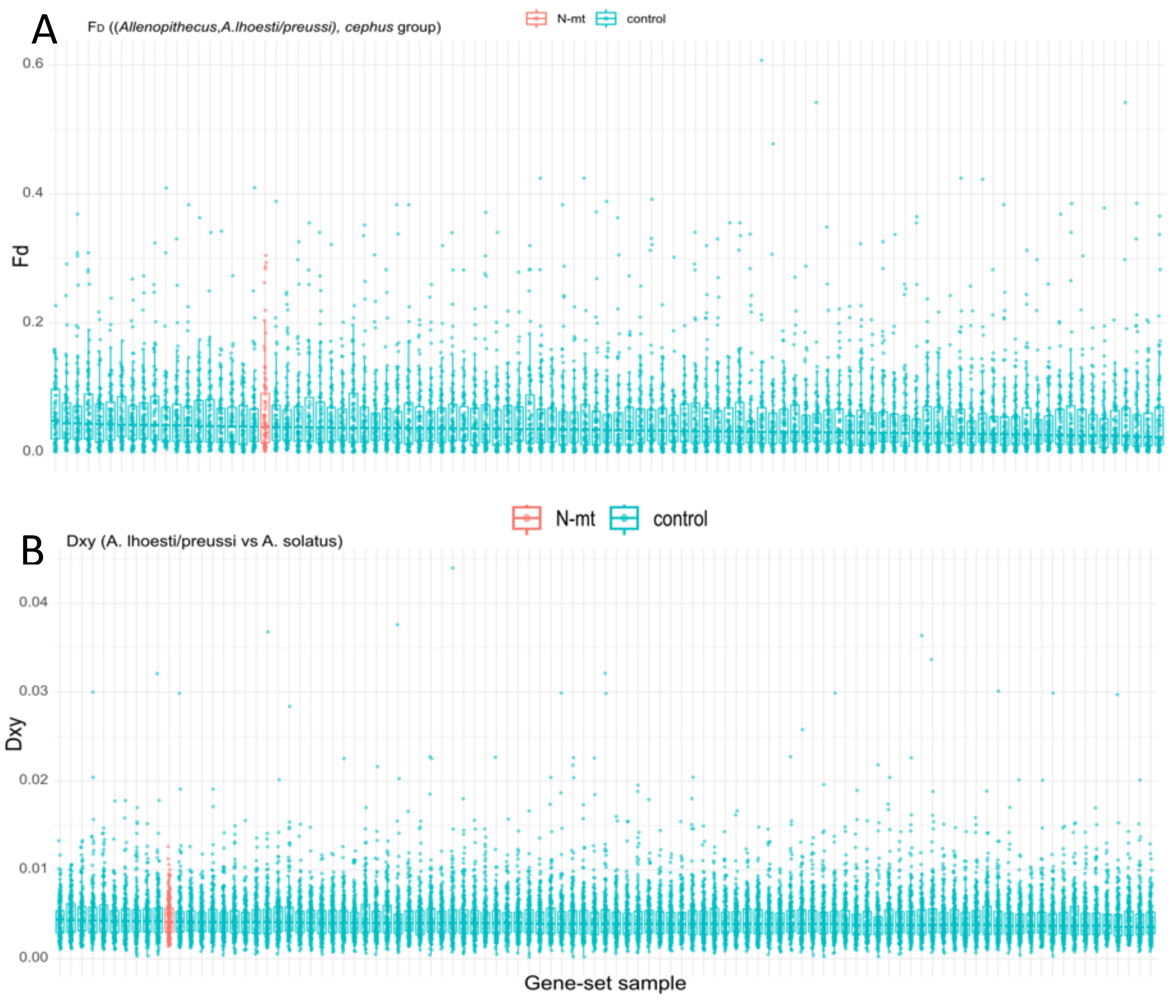


**Figure S25**. **Lack of broad-scale co-introgression of N-mt genes with mitochondria from the *cephus* group into *A. lhoesti/preussi.*** A) Introgression ($F_D$) from the *cephus* group into *A. lhoesti/preussi* in 199 nuclear genes that interact with the mitochondrion (N-mt, red) and 100 random samples of 199 genes that do not (green). These results show that the introgression levels are similar in N-mt compared to control genes. Genes with fewer than 200 SNPs were excluded. Gene sets are ordered by decreasing median Fd. B) Nucleotide divergence ($D_{XY}$) between *Allochrocebus lhoesti/preussi* and *A. solatus* in 199 nuclear genes that interact with the mitochondrion (N-mt, red) and 100 random samples of 199 genes that do not (green). These results show that the divergence is similar in the two groups of genes. Genes with fewer than 1,000 genotyped sites were excluded. Gene sets are ordered by decreasing median $D_{XY}$.

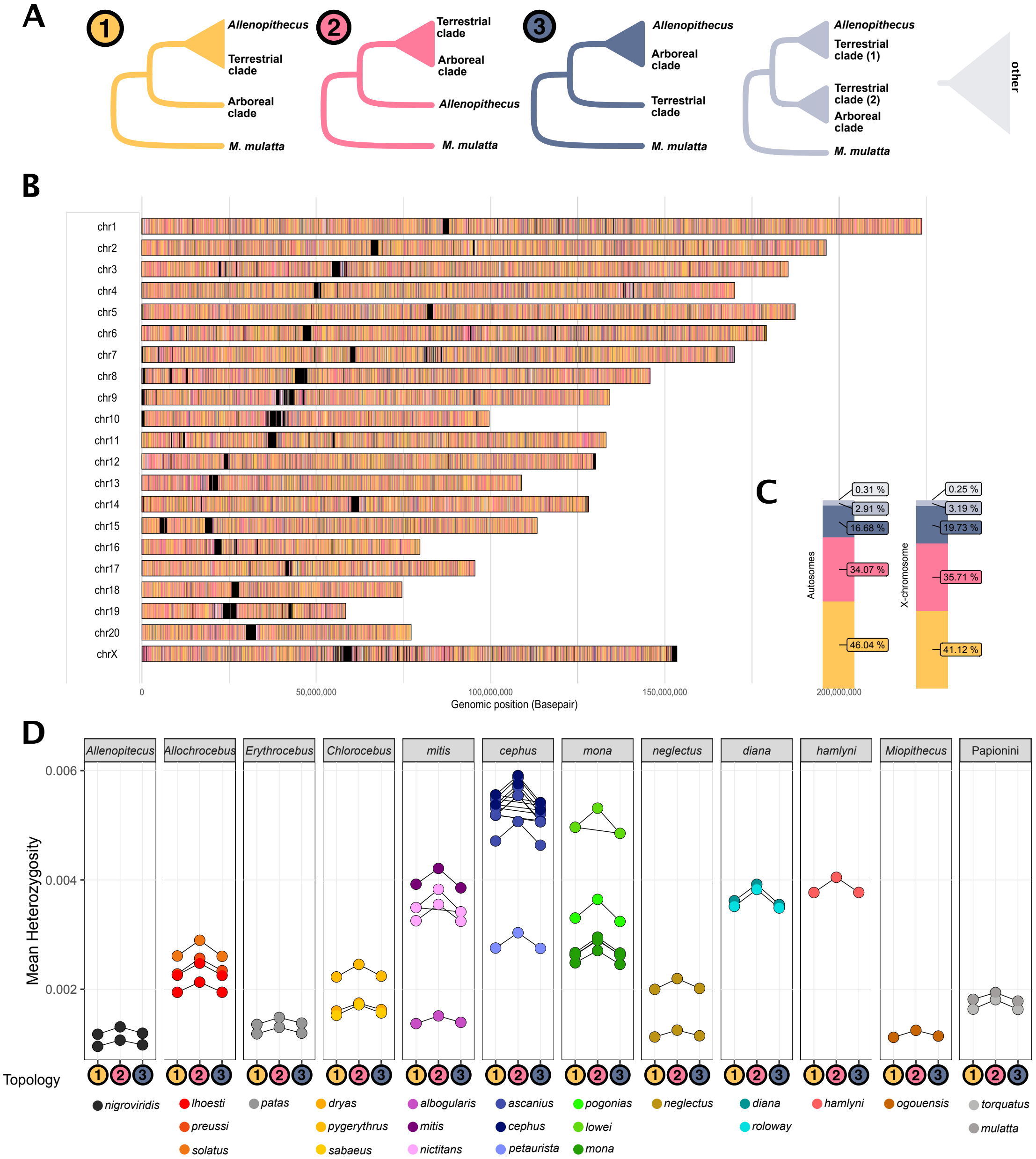


**Figure S26**. **Distribution, prevalence and heterozygosity of introgressed genomic segments in gene flow event B.** A) Five possible tree topologies obtained from 25 kb genomic windows for the introgression event B (Figure 3), including *Allenopithecus,* the Terrestrial clade and the Arboreal clade, rooted with *M. mulatta* as outgroup. Tree 1 shows the species tree. Tree 2 can arise through introgression or ILS. Tree 3 is expected only from ILS. Trees 4 and 5 show more complicated/unresolved topologies that can arise through e.g. more recent introgression or ILS. B) The genomic location of the different tree topologies along the *M. mulatta* chromosomes, and C) their relative abundance on the autosomes and the X-chromosome. Black blocks in (B) correspond to regions of the genome with insufficient information for inferences, frequently located around centromeres and telomeres. D) Heterozygosity per sample in genomic windows showing topology 1-3, connectors depict differences that were inferred as stastically significant in ANOVA tests with post hoc Tukey's test, corrected for multiple testing.

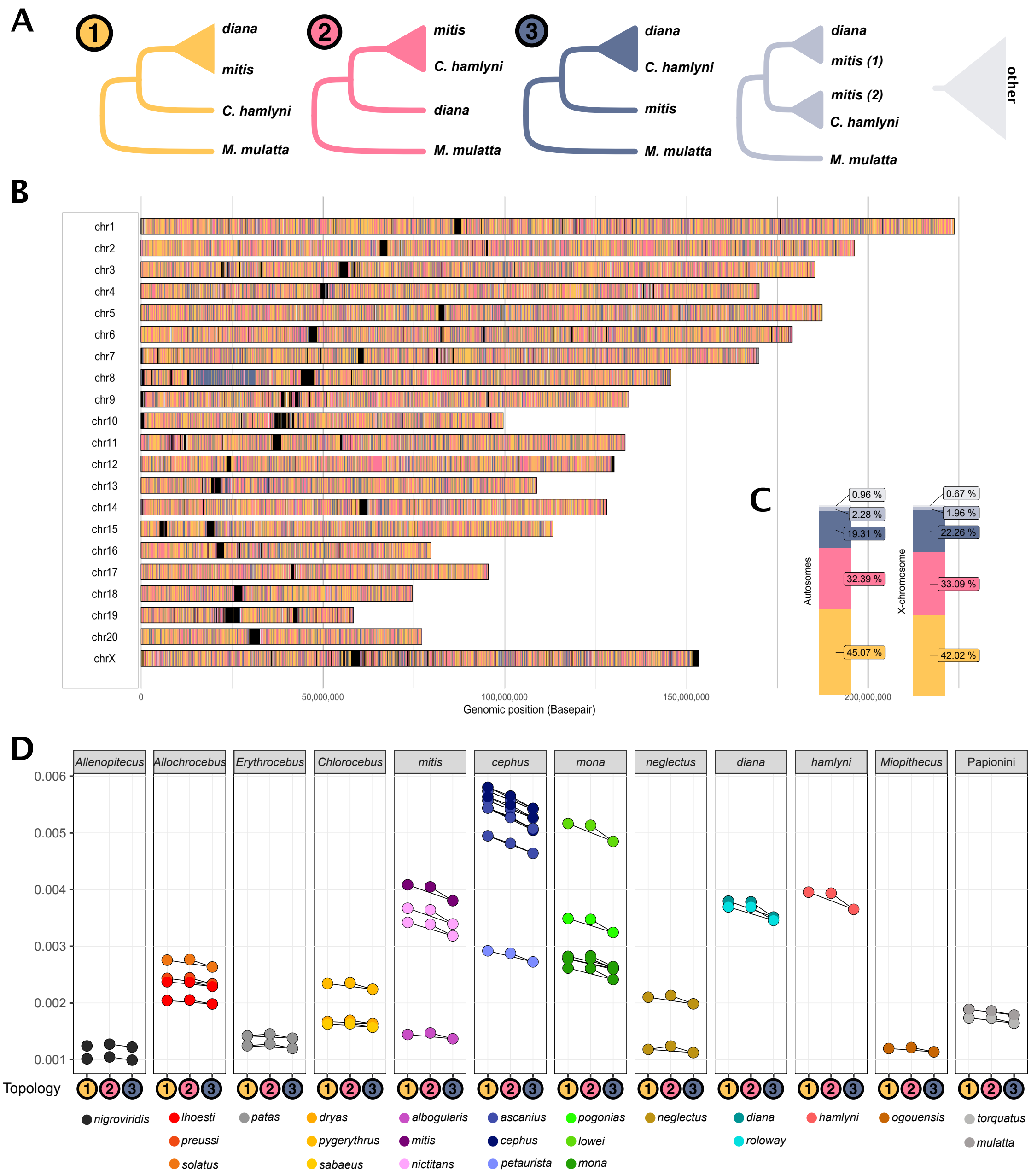


**Figure S27**. **Distribution, prevalence and heterozygosity of introgressed genomic segments in gene flow event C.** A) Five possible tree topologies obtained from 25 kb genomic windows for the introgression event C (Figure 3), including *C. hamlyni*, *mitis* and *diana* species groups, rooted with *M. mulatta* as outgroup. Tree 1 shows the species tree. Tree 2 can arise through introgression or ILS. Tree 3 is expected only from ILS. Trees 4 and 5 show more complicated/unresolved topologies that can arise through e.g. more recent introgression or ILS. B) The genomic location of the different tree topologies along the *M. mulatta* chromosomes, and C) their relative abundance on the autosomes and the X-chromosome. Black blocks in (B) correspond to regions of the genome with insufficient information for inferences, frequently located around centromeres and telomeres. D) Heterozygosity per sample in genomic windows showing topology 1-3, connectors depict differences that were inferred as stastically significant in ANOVA tests with post hoc Tukey's test, corrected for multiple testing.

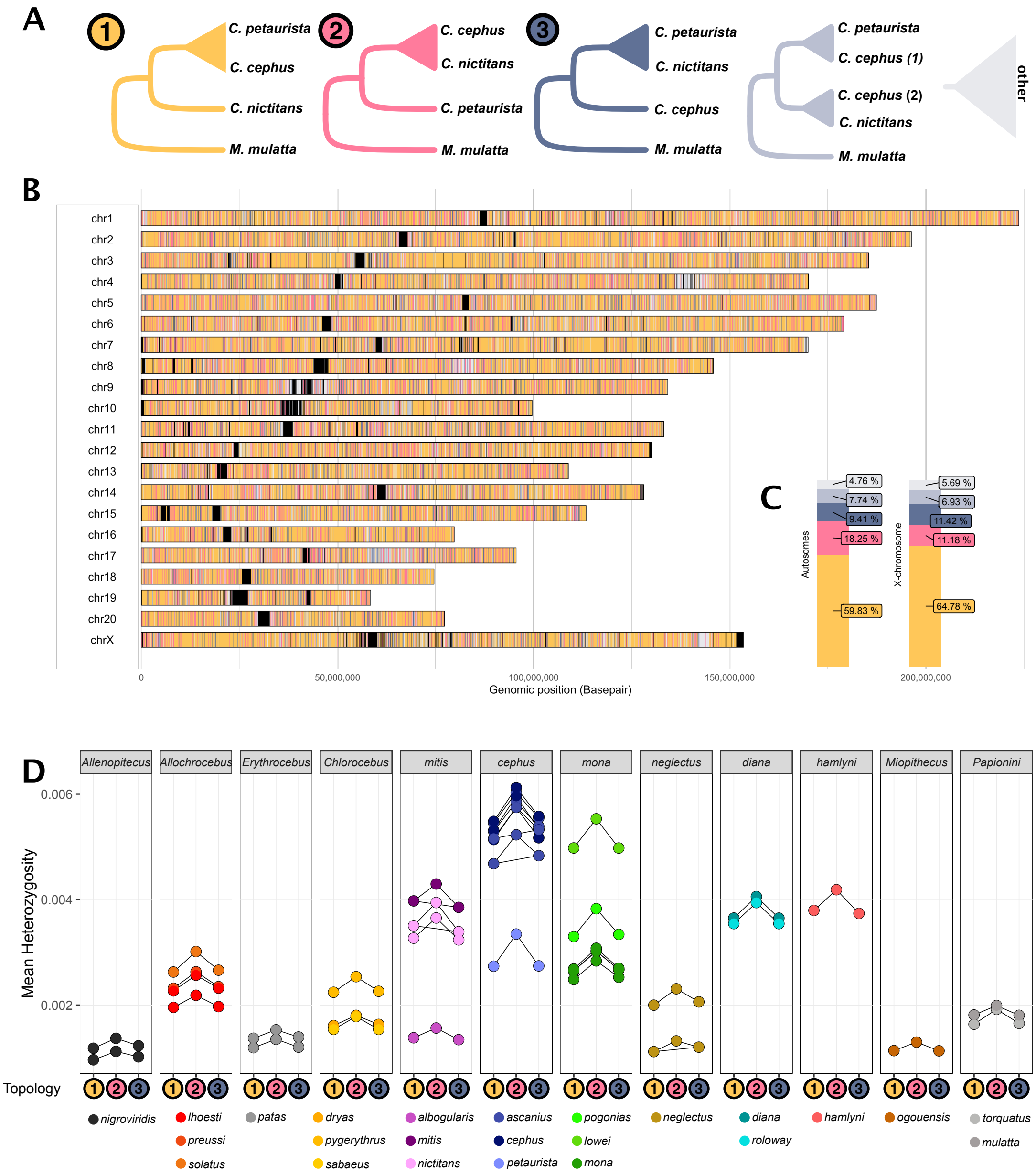


**Figure S28**. **Distribution, prevalence and heterozygosity of introgressed genomic segments in gene flow event D.** A) Five possible tree topologies obtained from 25 kb genomic windows for the introgression event D (Figure 3), including *C. petaurista, C. cephus/ascanius,* and the *mitis* group, rooted with *M. mulatta* as outgroup. Tree 1 shows the species tree. Tree 2 can arise through introgression or ILS. Tree 3 is expected only from ILS. Trees 4 and 5 show more complicated/unresolved topologies that can arise through e.g. more recent introgression or ILS. B) The genomic location of the different tree topologies along the *M. mulatta* chromosomes, and C) their relative abundance on the autosomes and the X-chromosome. Black blocks in (B) correspond to regions of the genome with insufficient information for inferences, frequently located around centromeres and telomeres. D) Heterozygosity per sample in genomic windows showing topology 1-3, connectors depict differences that were inferred as stastically significant in ANOVA tests with post hoc Tukey’s test, corrected for multiple testing.

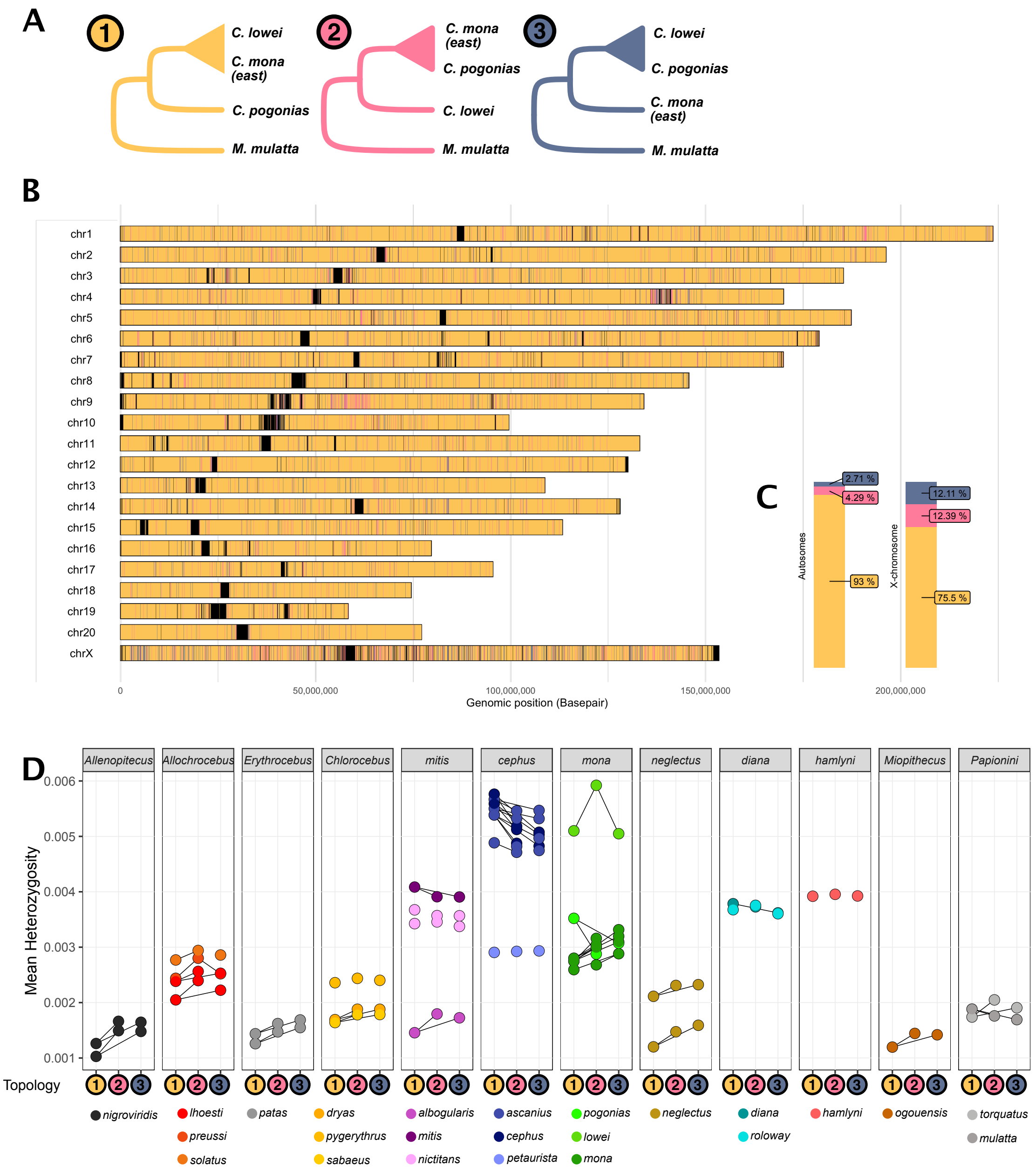


**Figure S29**. **Distribution, prevalence and heterozygosity of introgressed genomic segments in gene flow event E.** A) Five possible tree topologies obtained from 25 kb genomic windows for the introgression event D (Figure 3), including *C. lowei,* the easter *C. mona* population, and *C. pogonias*, rooted with *M. mulatta* as outgroup. Tree 1 shows the species tree. Tree 2 can arise through introgression or ILS. Tree 3 is expected only from ILS. Note that the two additional tree classes included in figure S27-29 [tree 4-5] are not applicable here because P2 and P3 consist of single samples. B) The genomic location of the different tree topologies along the *M. mulatta* chromosomes, and C) their relative abundance on the autosomes and the X-chromosome. Black blocks in (B) correspond to regions of the genome with insufficient information for inferences, frequently located around centromeres and telomeres. D) Heterozygosity per sample in genomic windows showing topology 1-3, connectors depict differences that were inferred as stastically significant in ANOVA tests with post hoc Tukey's test, corrected for multiple testing.

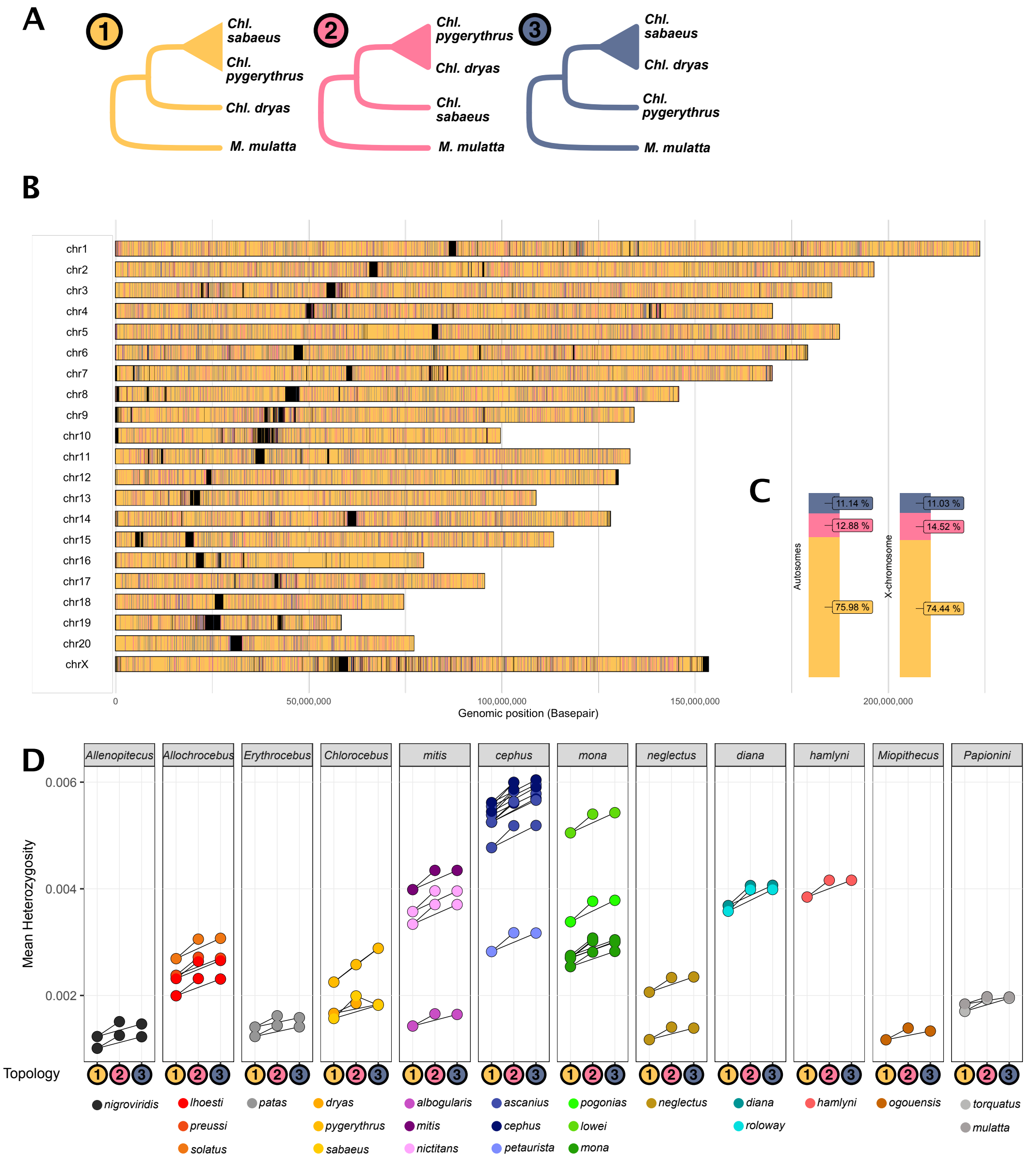


**Figure S30**. **Distribution, prevalence and heterozygosity of introgressed genomic segments in gene flow event F.** A) Five possible tree topologies obtained from 25 kb genomic windows for the introgression event D (Figure 3), including *Chl. sabaeus,* the easter *Chl. pygerythrus* population, and *Chl. dryas*, rooted with *M. mulatta* as outgroup. Tree 1 shows the species tree. Tree 2 can arise through introgression or ILS. Tree 3 is expected only from ILS. Note that the two additional tree classes included in figure S27-29 [tree 4-5] are not applicable here because P2 and P3 consist of single samples. B) The genomic location of the different tree topologies along the *M. mulatta* chromosomes, and C) their relative abundance on the autosomes and the X-chromosome. Black blocks in (B) correspond to regions of the genome with insufficient information for inferences, frequently located around centromeres and telomeres. D) Heterozygosity per sample in genomic windows showing topology 1-3, connectors depict differences that were inferred as stastically significant in ANOVA tests with post hoc Tukey's test, corrected for multiple testing.

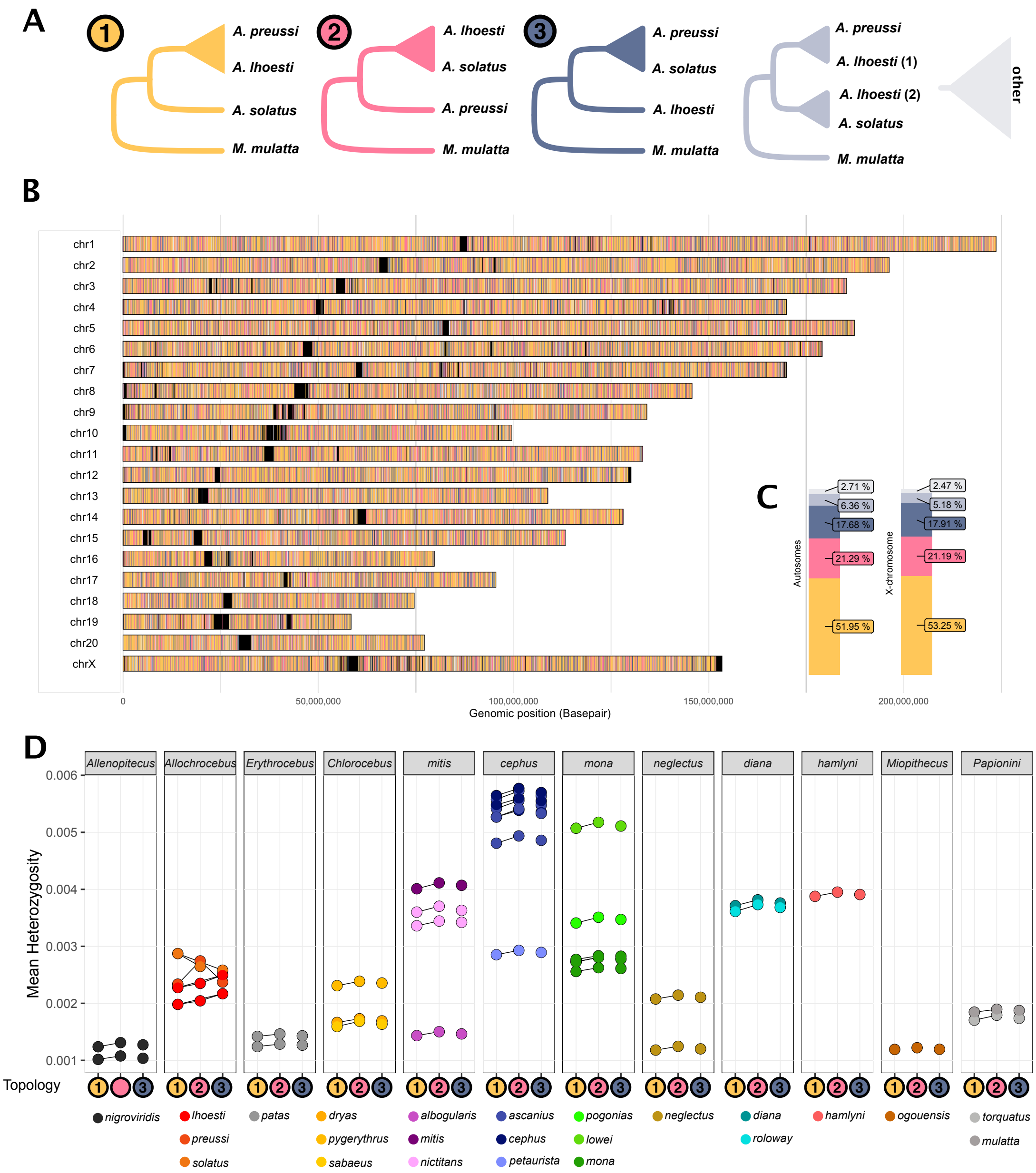


**Figure S31**. **Distribution, prevalence and heterozygosity of introgressed genomic segments in gene flow event G.** A) Five possible tree topologies obtained from 25 kb genomic windows for the introgression event D (Figure 3), including *A. preussi, A. lhoesti* and *A. solatus*, rooted with *M. mulatta* as outgroup. Tree 1 shows the species tree. Tree 2 can arise through introgression or ILS. Tree 3 is expected only from ILS. Trees 4 and 5 show more complicated/unresolved topologies that can arise through e.g. more recent introgression or ILS. B) The genomic location of the different tree topologies along the *M. mulatta* chromosomes, and C) their relative abundance on the autosomes and the X-chromosome. Black blocks in (B) correspond to regions of the genome with insufficient information for inferences, frequently located around centromeres and telomeres. D) Heterozygosity per sample in genomic windows showing topology 1-3, connectors depict differences that were inferred as stastically significant in ANOVA tests with post hoc Tukey’s test, corrected for multiple testing.

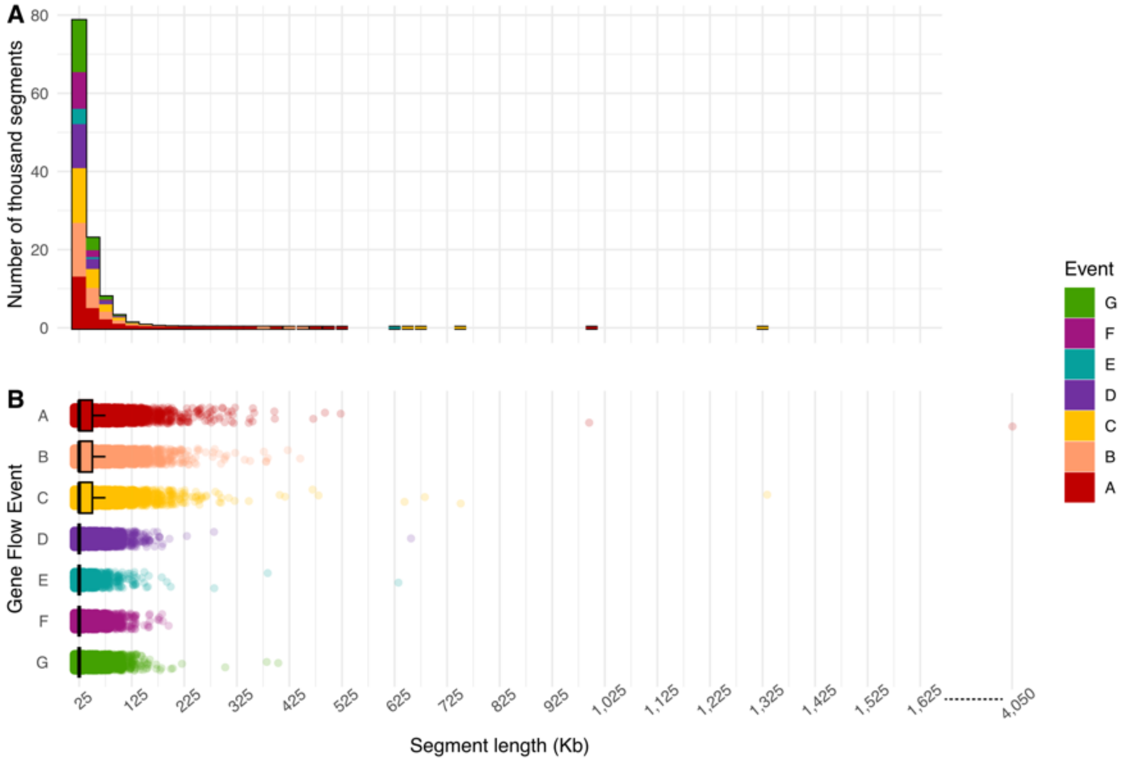


**Figure S32**. **Genomic regions of introgressed ancestry are short, in line with ancient introgression.** A) Counts of consecutive genomic regions of different lengths with ancestry in line with introgression (Tree 2) across all events, and B) the distributions of such regions. The vast majority of regions of introgressed ancestry were ≤ 25 kb (a single window), and long regions ≥ 100 kb were rare. This is in line with ancient timing of introgression, allowing recombination time to break down introgressed haplotype blocks.

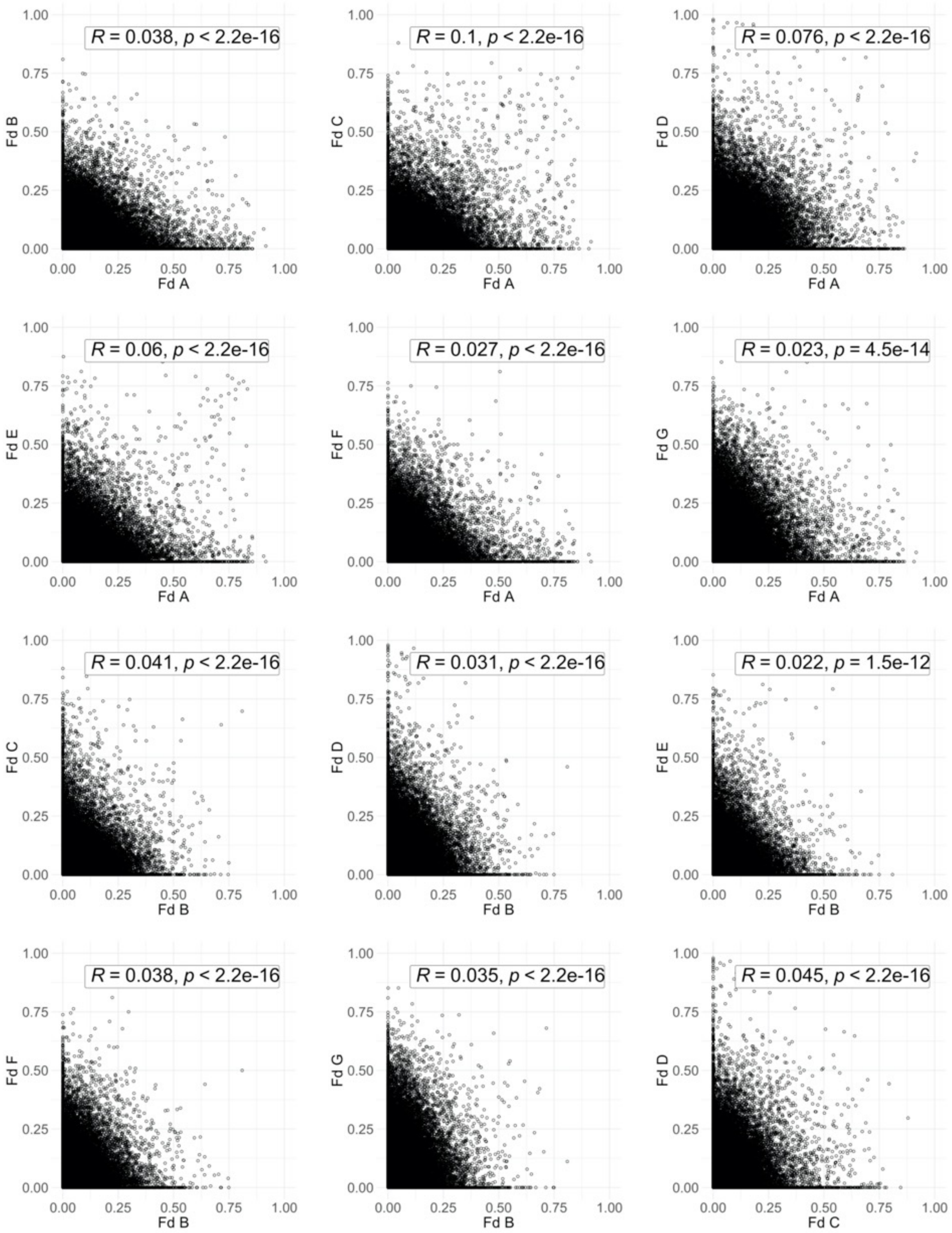


**Figure S33. Continuation and legend on next page**

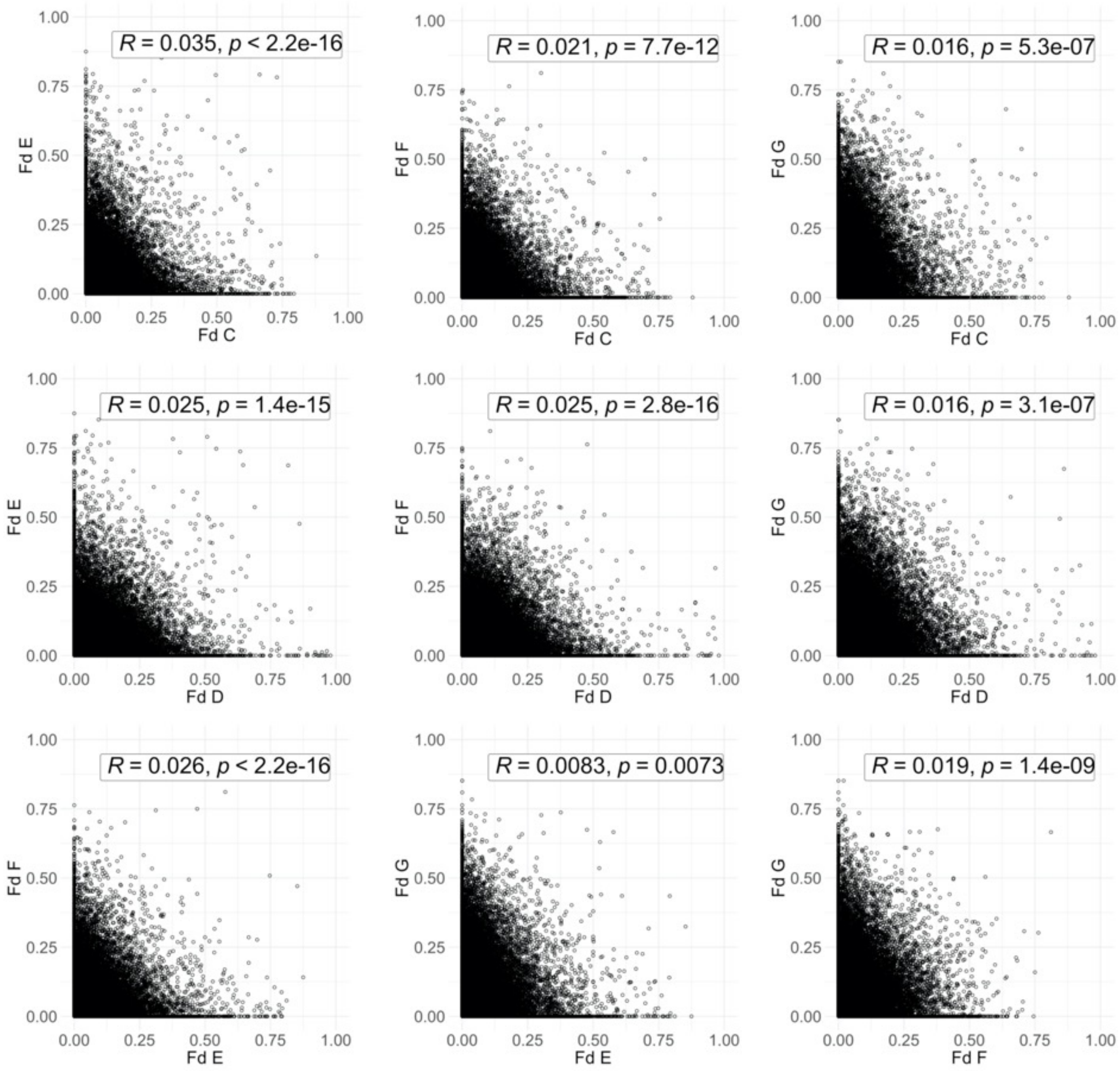


**Figure S33.** Pearsons correlation of Fd values in non-overlapping 25 kb windows along the reference genome, for all pairwise comparisons of independent gene flow events. Compared events are given on X and Y axis lables, each dot represent one 25 kb window.

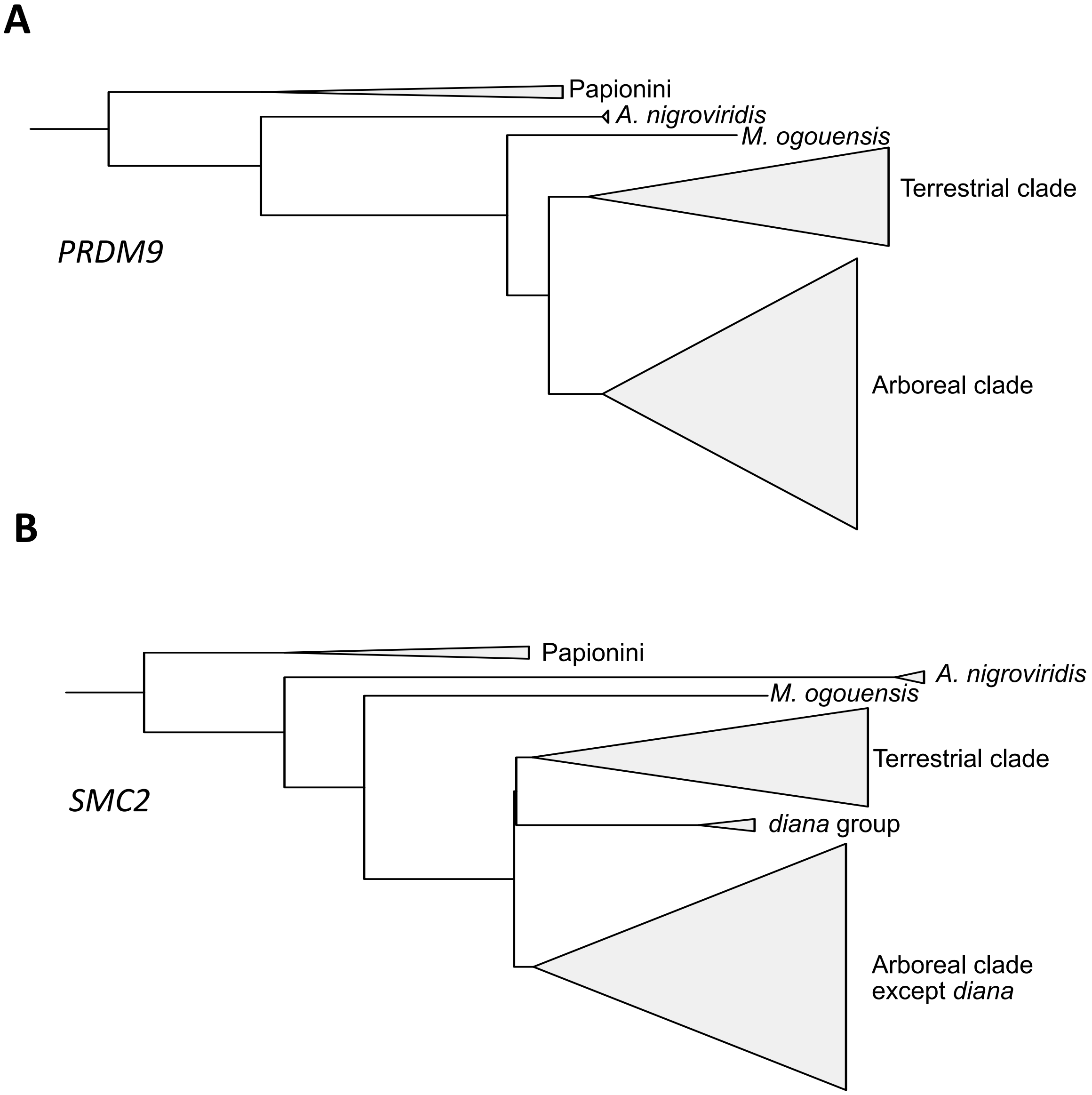


**Figure S34**. Neighbor joining trees in the 25 kb regions identified as introgressed between the Terrestrail and Arboreal clade ancestors (event B) that overlap the genes *PRDM9* (A) and *SMC2* (B). Note that these trees both suggest a sister relationship between the Terrestrial and Arboreal clades, and a closer relationship between these groups and *M. ogouensis* compared to *A. nigroviridis*. This suggests that introgression most likely occurred from the Arboreal clade ancestor into the Terrestrial clade, as the opposite direction would be expected to generate a closer relationship between these groups and *A. nigroviridis*.

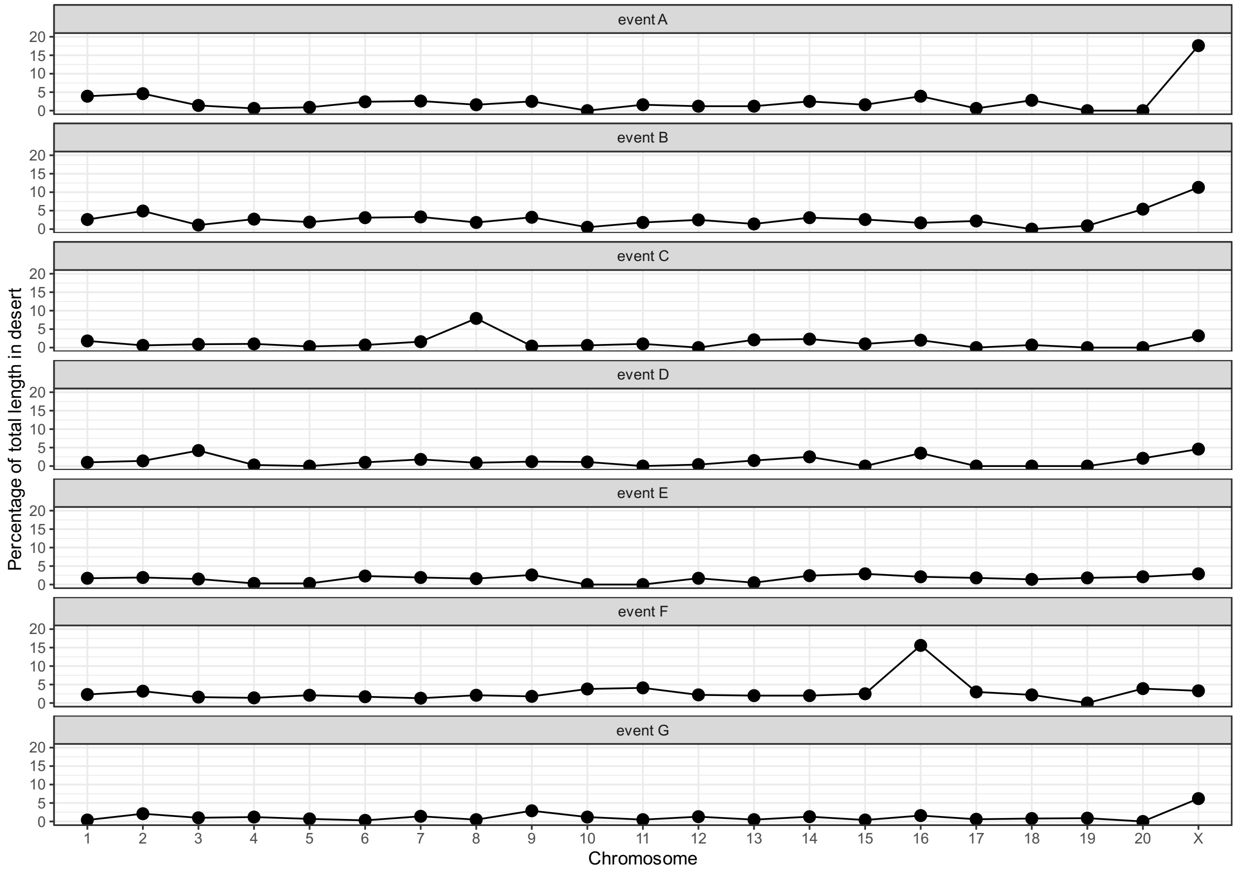


**Figure S35**. Proportion of autosomes (chromosomes 1-20) and the X-chromosome contained in introgression deserts, for each event.
